# The selection for cidofovir-resistant mutant vaccinia viruses is inhibited by coinfecting cidofovir-sensitive wildtypes

**DOI:** 10.64898/2026.09.14.751616

**Authors:** Stephen Z. Lee, Yi-Chan Lin, David H. Evans, Ryan S. Noyce

**Author notes:** Co-senior authors (David H. Evans, Ryan S. Noyce). Corresponding Authors: Ryan S. Noyce, 6-142N Katz Group Centre for Research, University of Alberta, Edmonton, AB, T6G 2R3, Canada., David H. Evans, 6-020 Katz Group Centre for Research, University of Alberta, Edmonton, AB, T6G 2R3, Canada.

## Abstract

Recent years have seen two outbreaks of zoonotic mpox, highlighting the need for additional therapies to treat these infections and curb the spread of the virus. However, there are only two antiviral drugs available for treating mpox, and drug resistance is a potential threat to intervention. Here we investigate how drug resistance emerges in a population of viruses where the sensitive wildtype strain vastly outnumbers the drug-resistant mutant. Using cidofovir treatment and resistance as a model system during vaccinia virus infection, we found that cidofovir-resistant poxviruses rapidly outcompete cidofovir-sensitive viruses when the two strains replicate in complete isolation. However, when the two strains infect the same cell, the drug-resistant virus lost its competitive advantage over the sensitive strain. We found that when the two viruses replicate in separate cells treated with cidofovir, the resistant strain displays faster genome replication kinetics compared to the sensitive strain. However, when the two viruses replicate in the same drug-treated cell, the strains replicated at the same rate. This showed that the drug-resistance trait was functionally shared between the two genetically-distinct strains, allowing the sensitive strain to parasitize the growth advantage of the resistant strain and preventing the rapid outgrowth of the resistant strain. These observations suggest that poxvirus coinfections negatively affect the selection for a rare trait like cidofovir-resistance. Considering that poxviruses actively prevent superinfection, we hypothesize that superinfection exclusion could maintain fitness in a population and facilitate the outgrowth of rare, advantageous traits in poxvirus populations.

**Author Summary:** Poxvirus infections remain a threat to public health. Treating these infections with a single therapeutic, while most often the only option, is ultimately not an ideal strategy as it is much easier for the virus to adapt to a single drug. Ideally, there would be multiple drugs used in combination to treat poxvirus infections as it is much harder for the virus to adapt to all the drugs in the cocktail without losing its ability to replicate. Our data support this paradigm, showing that a poxvirus population with resistant mutants can adapt to a single antiviral drug within one round of replication. These data agree with others in the field that have shown that poxviruses can very rapidly adapt to selection from antiviral drugs. This stems from multiple molecular strategies that poxviruses employ to adapt to selective pressures. As more and more outbreaks of poxvirus infection occur, we need to be prepared to deploy multiple treatment strategies to curb the spread of the pathogen.

## Introduction

Poxviruses are remarkably successful pathogens. Co-evolution with their hosts has produced an arsenal of strategies to rapid adaptation to selective pressures, either from the host immune system or other forms of selection^1,2^. One such form of selective pressure is drug treatment. The global outbreak of monkeypox virus (MPXV) highlighted the need for effective therapeutics to treat infection and increased the use of three existing anti-poxvirus drugs in humans^3,4^. Tembexa (brincidofovir, BDV), and Vistide (cidofovir, CDV) are the only drugs licensed for treating Orthopoxvirus infections. Tecovirimat inhibits the viral phospholipase, which in vaccinia virus (VACV) is the F13 protein^5^. BDV is a prodrug of CDV, an acyclic nucleoside phosphonate analogous to deoxycytidine triphosphate (dCTP) that inhibits the DNA polymerase E9^6–9^. Seeing as these drugs are only approved as monotherapies, there is concern that drug resistance can rapidly arise, rendering treatment ineffective. Tecovirimat-resistance is well-documented in circulating MPXV isolates^10,11^. Conversely, there is less known about the ability of CDV resistance (CDV^R^) to evolve in poxvirus populations.

*In vitro*, CDV acts as a delayed chain terminator, with synthesis stopping at the n+1 position of the daughter strand^8^. CDV also exerts a template-dependent effect, where the polymerase fails to add dGTP across the CDV lesion^7^. CDV^R^ arises from substitutions in the E9 at positions A314 and A684^12,13^. Position 314 is in the exonuclease domain, and an A314T substitution enhances the excision of CDV from the daughter strand^14–17^. Position 684 is adjacent to the nucleotide binding pocket, and an A684V substitution is thought to prevent CDV entry into the active site^13^. These substitutions arose after nearly 40 passages of the virus with increasing concentrations of CDV^12^. This virus is not attenuated in tissue culture but is attenuated *in vivo*. It is striking that, despite there being no fitness cost during passage *in vitro*, selection for CDV^R^ required such extensive passage. This is especially interesting seeing how selection for other drug resistance mutations can occur after a single passage^5,18–20^. We therefore wanted to know how the selection for CDV^R^ operates in the population.

Generally, if individual viruses were replicating in total isolation, then the CDV-sensitive (CDV^S^) wildtypes should not survive after drug treatment, while the CDV^R^ viruses do. Thus, one would expect rapid outgrowth of the CDV^R^ virus. However, it becomes unclear how the selection would operate if CDV^R^ and CDV^S^ viruses coinfected cells. The poxvirus lifecycle occurs in cytoplasmic factories that form early during infection from DNA binding proteins interacting with the uncoated genome and the host endoplasmic reticulum^21–23^. DNA replication, late gene expression, translation, and assembly occur in or around the factory^24–28^. We previously demonstrated that the virus factories affect interactions between coinfecting VACVs, physically separating the genomes and preventing inter-virus recombination^29–31^. Additionally, there is evidence that the factory can sequester newly synthesized proteins^24,31^. We previously used recombinant viruses that expressed the λ bacteriophage cro DNA binding domain fused to enhanced green fluorescent protein (EGFP) or monomeric Cherry fluorescent protein (mCherry)^31^. Coinfection with these viruses generated factories with either EGFP-cro or mCherry-cro fluorescence, but not both.

If the factory can sequester a DNA binding protein, then one would expect that in a coinfection, the CDV^R^ polymerase would only replicate the CDV^R^ genome, allowing rapid outgrowth of the mutant despite the presence of CDV^S^ viruses. Why then was selection for CDV^R^ so slow? We sought to reconcile these differences by conducting coinfection experiments with CDV^R^ and CDV^S^ viruses. Surprisingly, we found that the CDV^S^ virus parasitized the growth advantage of the CDV^R^ virus during a coinfection in CDV-treated cells, allowing the CDV^S^ virus to survive at a high titer in the population. When the two strains replicated in isolation, there was rapid purifying selection for the CDV^R^ virus with drug treatment. These results suggest that coinfections between strains of differing fitness can negatively affect the population. Additionally, these results suggest that the selection for a CDV^R^ mutant could occur very rapidly during drug treatment, complicating the treatment of Orthopoxvirus infections in patients.

## Results

### Construction of recombinant VACVs expressing cro-EGFP and cro-mKate2 proteins

Previously, our laboratory used viruses expressing EGFP-cro and mCherry-cro to track VACV factories using live-cell imaging^31,32^. However, expression of the reporter attenuated the growth of the viruses in cell culture. Therefore, we constructed a second group of recombinant VACVs that expressed cro-EGFP or cro-mKate2 (Figure 1A). We used VACV Western Reserve as the CDV^S^ strain and VACV VDG1.3 as the CDV^R^ strain. The cro-EGFP and cro-mKate2 proteins allowed us to differentiate between virus plaques in a monolayer and to track virus factories by live-cell imaging (Figure 1B and C). Immunofluorescence using a monoclonal antibody targeting the VACV single-stranded DNA binding protein I3 confirmed that cro-EGFP and cro-mKate2 specifically labeled virus factories (Figure 1D). Additionally, we conducted multi-step growth curves to assess the fitness of the recombinant viruses relative to the parental strains. We found that expression of cro-EGFP and cro-mKate2 did not significantly reduce virus fitness, as we found no difference in the infectious titers between the recombinant viruses and the parental strains after 48 hours of replication (Figure S1A). Lastly, we conducted plaque reduction assays to determine the half maximal effective concentration (EC_50_) of CDV. As expected, the viruses expressing the CDV^R^ polymerase were significantly less sensitive to CDV as there was a significant increase in EC_50_ for the CDV^R^ strains compared to the corresponding CDV^S^ strains (Figure S1B).

**Figure 1.**
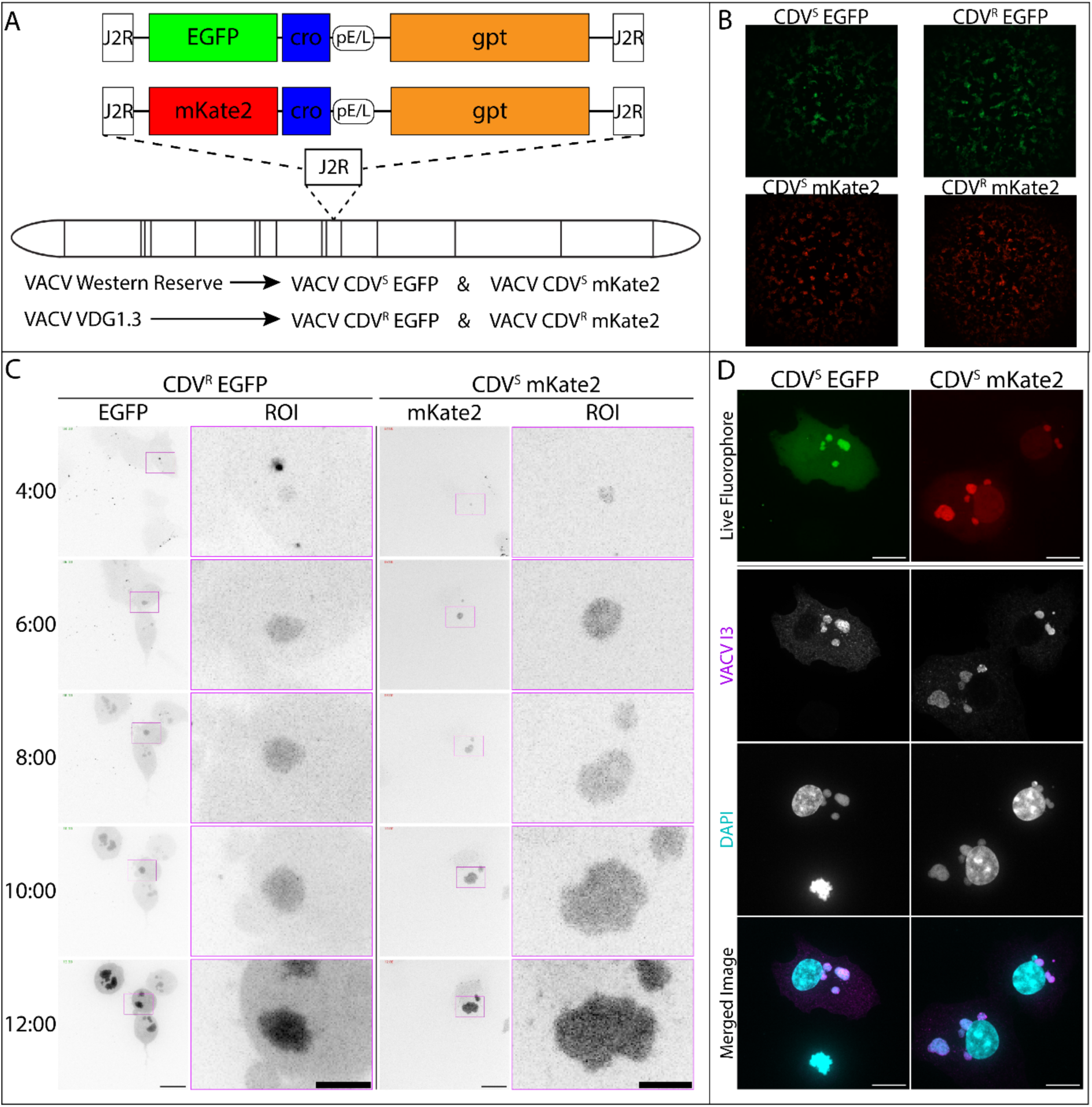
Generation of viruses expressing cro-EGFP and cro-mKate2. A) Schematic representation showing the insertion of cro-EGFP or cro-mKate2 into the VACV *J2R* gene. B) Virus plaques imaged at 48 hours post-infection. BSC-40 cells were infected with the indicated VACV strains and imaged using an EVOS FL Auto inverted microscope. C) Live-cell imaging showing cro-EGFP and cro-mKate2 distribution during virus replication. BSC-40 cells were infected with the indicated VACV strains, and the cells were imaged from 1-12 hours post-infection using a spinning disk confocal microscope. Scale bar = 15µm, region of interest (ROI) scale bar = 5µm. D) cro-EGFP and cro-mKate2 label virus factories. BSC-40 cells growing in gridded live-cell dishes were infected with the indicated VACV strains and imaged at 6 hours post-infection. Virus factories were visualized using an anti-I3 antibody and DAPI staining. The live-cell grid was used to relocate the cells, which were re-imaged. Scale bars = 15µm.

### CDV^R^ viruses lose their competitive advantage over CDV^S^ viruses in coinfected cells

To examine CDV^R^ selection, we measured viral frequency after a single round of replication in the presence of CDV. We first used a low MOI (MOI 0.02) to model how the population changed when, on average, cells received a single virus particle (Figure 2A). We designated the diluted virus mixture as the Input and the progeny from CDV^+^ or CDV^-^ cells as Output CDV^+^ or CDV^-^, respectively. We re-plated each sample on fresh BSC-40 cells without CDV treatment to generate a mixture of plaques that allowed us to quantify the proportion of the two strains in the population. The ratio of viruses in the Input mixture showed a higher proportion of CDV^R^ viruses than anticipated (Figure 2C). However, the population randomly drifted back to 1:1 when replicating in CDV^-^ cells (Figure 2C). After replication in CDV^+^ cells, the CDV^R^ virus rapidly outcompeted the CDV^S^ strain, becoming dominant in the population (Figure 2C). Thus, when the CDV^R^ and CDV^S^ viruses replicate in complete isolation, purifying selection occurred and rapidly selected for the CDV^R^ virus. We also titrated the samples and found that the concentration of viruses at Input was approximately 10^4^ PFU/mL (Figure S2A). With our infection conditions, this would correspond to 3000 PFU delivered to each well, meaning there was a very low probability of cells receiving multiple particles. These experiments showed that a low MOI, ensuring the viruses replicated in isolation, allowed rapid adaptation and advanced population fitness.

**Figure 2.**
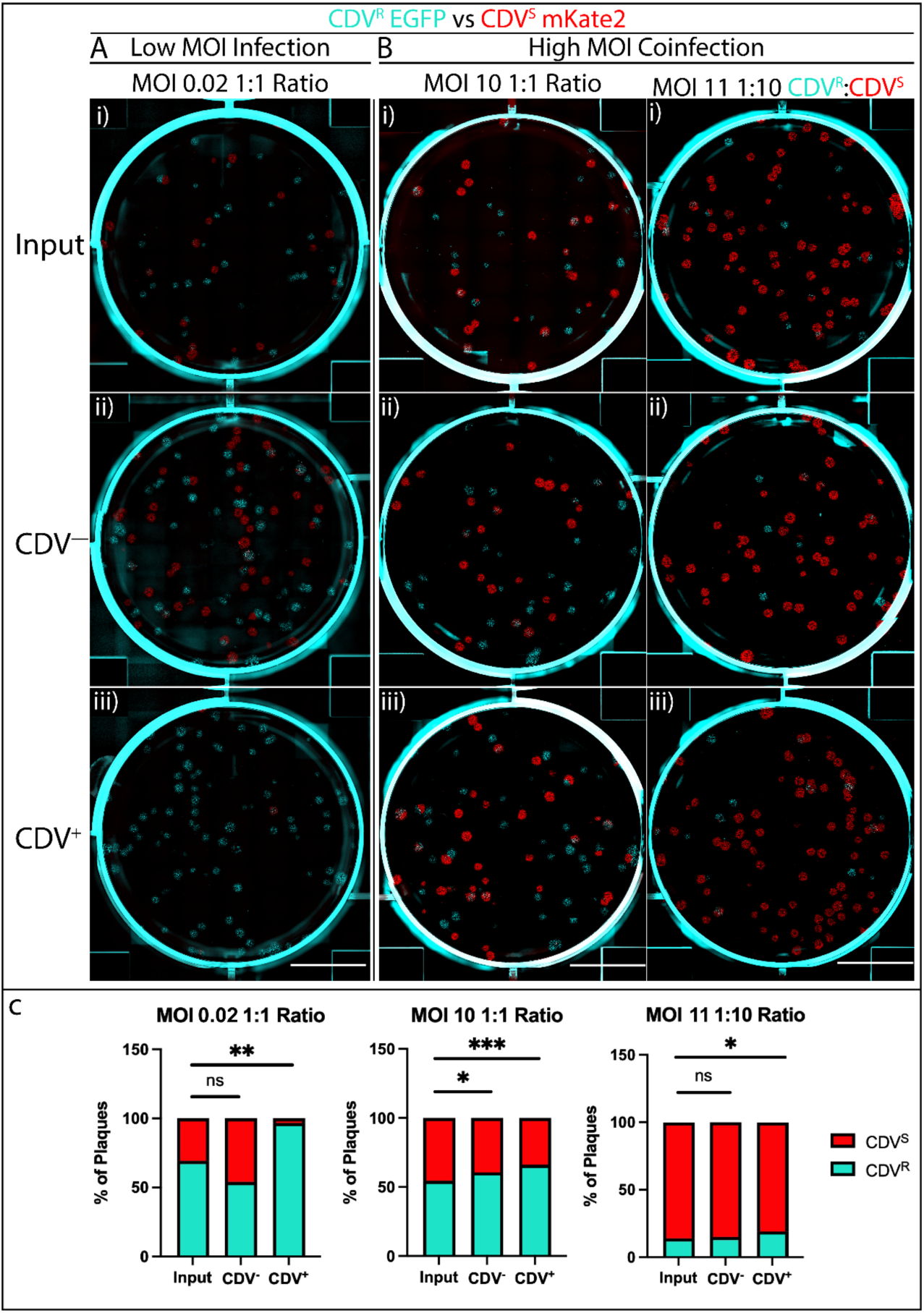
CDV^R^ VACV does not outcompete CDV^S^ VACV during a coinfection. Diluted virus (input) and progeny from the infections in the absence (CDV^--^) or presence (CDV^+^) of CDV were plated on fresh, untreated BSC-40 cells calculate the frequency of CDV^R^ (cyan) and CDV^S^ (red) viruses. Plaques were visualized 48 hours post-infection using a Biotek Cytation at 1.25X magnification. Representative wells from three independent biological experiments are shown. Scale bar = 1cm. A) CDV^R^ and CDV^S^ viruses were diluted to a combined MOI of 0.02 at a 1:1 ratio to model a scenario in which the two viruses replicate in isolation. B) A combined MOI of 10 with a 1:1 ratio of CDV^R^ and CDV^S^ viruses was used for a coinfection model. An MOI of 11 with a 1:10 ratio of CDV^R^ and CDV^S^ viruses was added to model a scenario in which a rare mutant has arisen in the population. C) The data from Figure 2 were quantified by counting red (CDV^S^) and cyan (CDV^R^) plaques and calculating the frequency of the two strains in the population. The data were analyzed with a two-way ANOVA with Dunnett’s multiple comparisons between the progeny from CDV^−^ and CDV^+^ cells and the Input. Values are the mean of three independent biological experiments.

Next, we conducted high-MOI coinfections using an MOI of 10 with a 1:1 ratio of CDV^R^:CDV^S^ viruses (Figure 2B). The concentration of the Input mixture was approximately 10^7^ PFU/mL, sufficiently high to ensure that the cells received multiple viruses (Figure S2B). Plating the progeny from CDV^-^ and CDV^+^ cells showed that the proportion of CDV^R^ viruses significantly increased in both conditions (Figure 2C). Random drift could explain these increases, and most notably, this experiment did not show any rapid outgrowth of the CDV^R^ strain as seen in Figure 2A. Additionally, we used an MOI of 11 with a 1:10 ratio of CDV^R^:CDV^S^ viruses to simulate a scenario early during the evolution of the trait, where this strain would be at a lower frequency within the population (Figure 2B). There was a significant increase in the proportion of CDV^R^ viruses in the progeny from CDV^+^ cells (Figure 2C). However, the CDV^R^ viruses comprised only 20% of the population. The vast majority of the CDV^S^ viruses survived the selection. These experiments showed that, surprisingly, coinfection negatively affected population fitness, allowing the less-fit CDV^S^ strain to survive the purifying selection from drug treatment and preventing the outgrowth of the more fit CDV^R^ strain.

### CDV^S^ virus factories grow slower than CDV^R^ factories under CDV selection

Based on the results from Figure 2, we wanted to investigate what was happening in single cells during a coinfection between CDV^R^ and CDV^S^ viruses. As shown in Figure 1, the expression of cro-EGFP allowed us to track virus factories using live-cell imaging. Because each virus factory forms from a single infecting genome^30^, we wanted to use this technique to track the CDV^S^ and CDV^R^ viruses with selection from CDV treatment. However, based on reviewing previous studies, it was unclear how CDV treatment would affect the growth of the virus factories. The biochemical and genetic evidence showing that mutations in the *E9L* gene promote CDV^R^ would suggest that CDV would inhibit factory growth as the rate of DNA replication decreases^7,8,12,13^. Conversely, experiments using CDV treatment in infected cells showed that CDV does not inhibit DNA replication, instead causing an encapsidation defect after the polymerase incorporates the drug into nascent DNA strands^33^. We sought to reconcile these conflicting reports and conducted mechanistic studies to determine if CDV would inhibit virus factory growth.

We used a pre-treatment strategy in which we added 330μM CDV to the cells overnight, then infected with VACV. This differed from previous studies in which the authors treated the cells with CDV concurrent with virus infection^33^. We reasoned that a pre-treatment would allow ample time for CDV conversion to CDVpp, which is a very slow process in cell culture^34^. In live-cell imaging, CDV^S^ virus factories showed rapid exponential growth from 4-6 hours post-infection in the absence of CDV selection (Figure 3A). These experiments show that the fluorescence intensity from cro-EGFP increased as viral DNA accumulated in the factories and gene expression continued (Figure 3A). Conversely, in cells treated with CDV, CDV^S^ virus factories formed by 4hpi but did not appreciably grow during the time course (Figure 3B). When we repeated these experiments using a CDV^R^ virus, there was a clear growth advantage from encoding the CDV^R^ polymerase, as the factories continued to replicate in the presence of CDV (Figure 3C and D).

**Figure 3.**
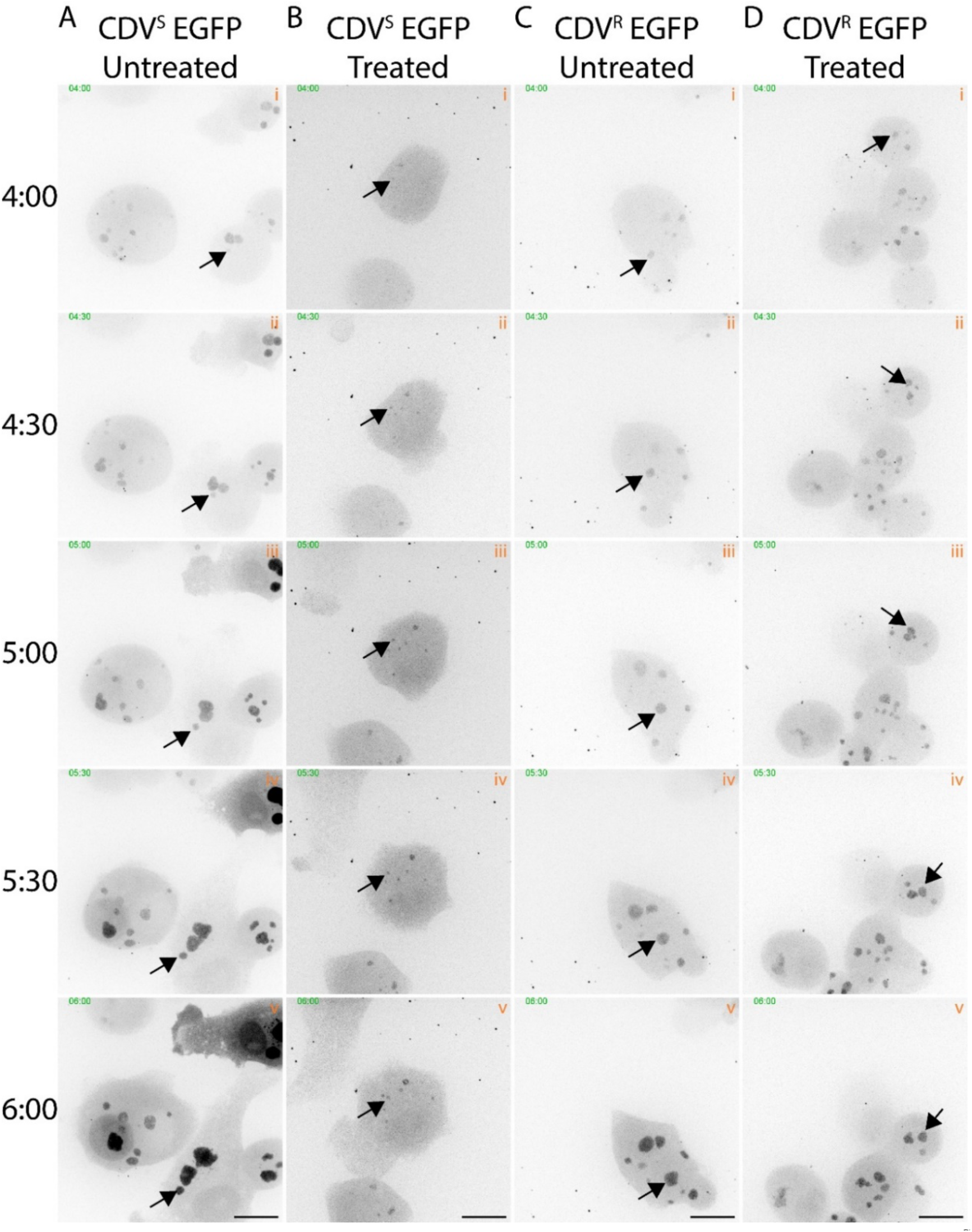
Live-cell imaging of VACV factories shows different growth kinetics of CDV^S^ and CDV^R^ viruses under CDV selection. BSC-40 cells were synchronously infected with VACV CDV^S^ EGFP (A & B) or VACV CDV^R^ EGFP (C & D) at an MOI of 5. Live-cell imaging was conducted from 3-6 hours post-infection by collecting 1µm z-stacks every 10 minutes. The z-stacks were merged based on fluorescence intensity and images were processed in Fiji. Scale bar = 15µm. Arrows were added to follow single factories in Adobe Illustrator during figure assembly.

To compare the growth of the two strains, we analyzed still frames from live-cell imaging, quantifying the size of factories and plotting these measurements as functions of time. Non-linear regression analysis showed that the CDV^S^ strain grew exponentially over time in untreated cells, but in CDV-treated cells, these factories displayed much flatter growth curves than the factories replicating in the absence of CDV (Figure 4A and B). Conversely, the CDV^R^ factories displayed strong exponential growth in the absence or presence of CDV, showing that the mutations in *E9L* allowed the virus to continuously replicate its genome and overcome the inhibitory effect of drug treatment (Figure 4C and D). To quantify this phenotypic difference, we extracted the doubling time from each growth curve and conducted statistical analysis (Figure 4E and F). On average, the doubling time for the CDV^S^ virus factories was significantly higher in CDV-treated cells compared to the untreated control (p<0.01). This showed that the growth rate of the CDV^S^ virus factories slowed down as the drug exerted an antiviral effect during DNA replication. There was no significant difference between the doubling times of the CDV^R^ factories replicating in the absence or presence of CDV. However, the doubling time for the CDV^R^ factories in treated cells was significantly lower than the CDV^S^ factories in this condition.

**Figure 4.**
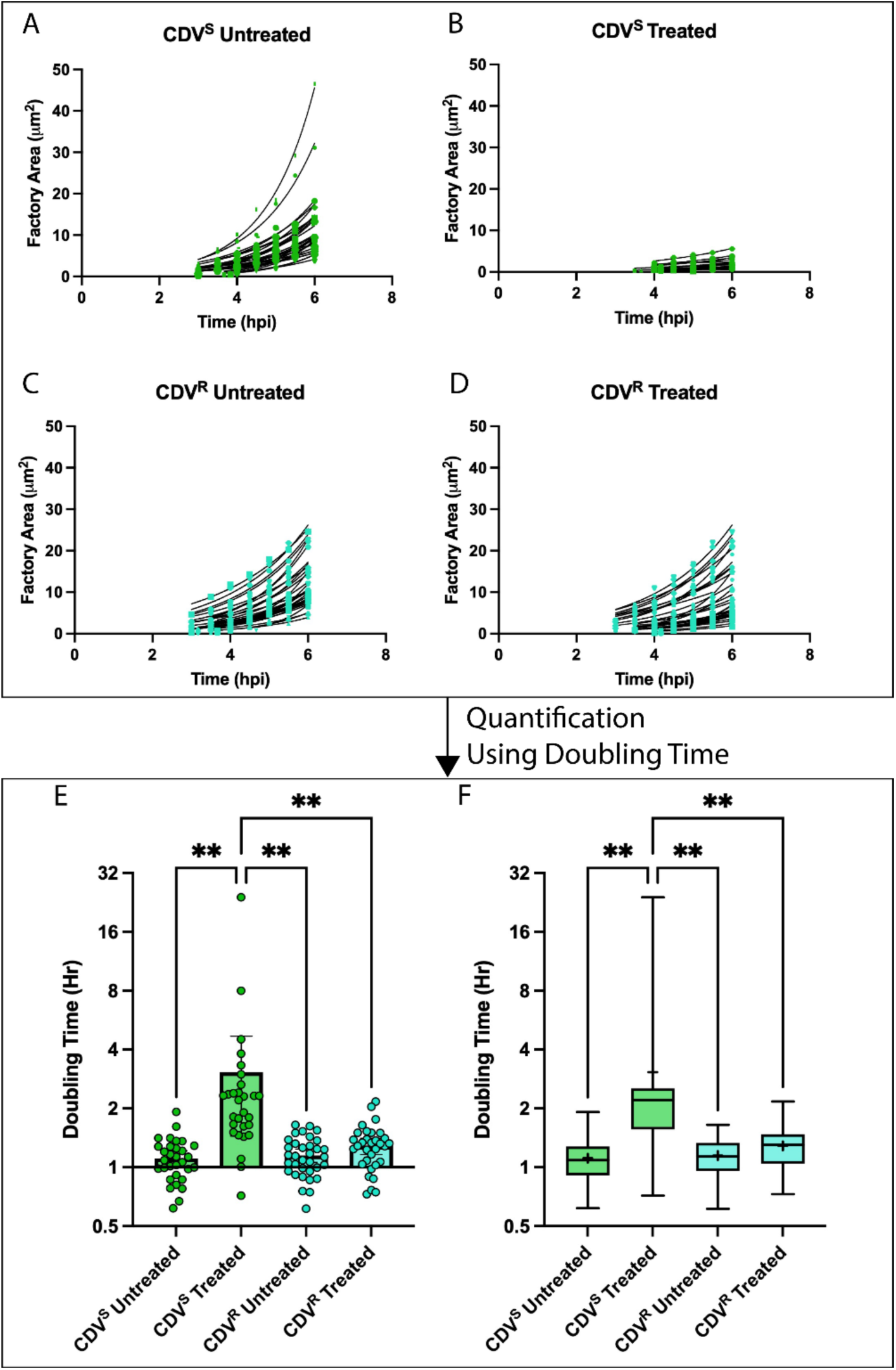
CDV^R^ virus factories have a growth advantage over CDV^S^ virus factories. A-D) Growth curves of virus factory growth. More than 30 virus factories imaged in Figure 4 were measured in FIJI from 3-6 hours post-infection. Only virus factories that did not collide with adjacent factories were measured in this analysis. Also, only virus factories that were continuously observed throughout the time course were included. E-F) The doubling times of the non-linear regression curves shown in A-D were compiled and analyzed using a one-way ANOVA with Tukey’s multiple comparisons. Mean and 95% confidence intervals are plotted. For E, + = mean, ̶ = median, box = first and third quartile, whisker = minimum and maximum.

### CDV inhibits viral DNA synthesis

To increase the number of virus factories and cells included in the analysis, we infected BSC-40 cells with CDV^R^ or CDV^S^ viruses in the absence or presence of CDV and fixed the samples at 6 hours post-infection. This also allowed us to include viruses expressing cro-mKate2 in the analysis that were not included in live-cell imaging experiments. Immunofluorescence microscopy showed that, in the absence of CDV, all four viruses assemble large factories in BSC-40 cells (Figure 5A and Supplemental Figure 3A). There was dense DAPI staining inside these factories, showing robust DNA replication (Figure 5A, panels i & iii). As expected, the I3 protein colocalized with DAPI. In CDV-treated cells, the CDV^S^ virus factories were much smaller than the CDV^R^ factories (Figure 5B and Supplemental Figure 3B). The CDV^S^ virus factories were punctate in the cell (Figure 5B, panels ii & iv), consistent with the phenotype observed in live-cell imaging (Figure 3B). Quantification of the DAPI fluorescence in each factory showed that in the absence of CDV, there were no statistical differences between the volume of CDV^S^ and CDV^R^ factories (Figure 5C). The expression of cro-EGFP or cro-mKate2 also did not affect factory size. However, with CDV pretreatment, the CDV^S^ factories were significantly smaller than the CDV^R^ factories (Figure 5C). By using DAPI fluorescence for quantification, this experiment also showed that there was significantly less DNA inside the CDV^S^ factories than the CDV^R^ factories (Figure 5B panels iii & v). This observation confirmed the data from Figure 4 showing that CDV does directly inhibit DNA synthesis and that the mutations in *E9L* allow the virus to replicate its genome normally.

**Figure 5.**
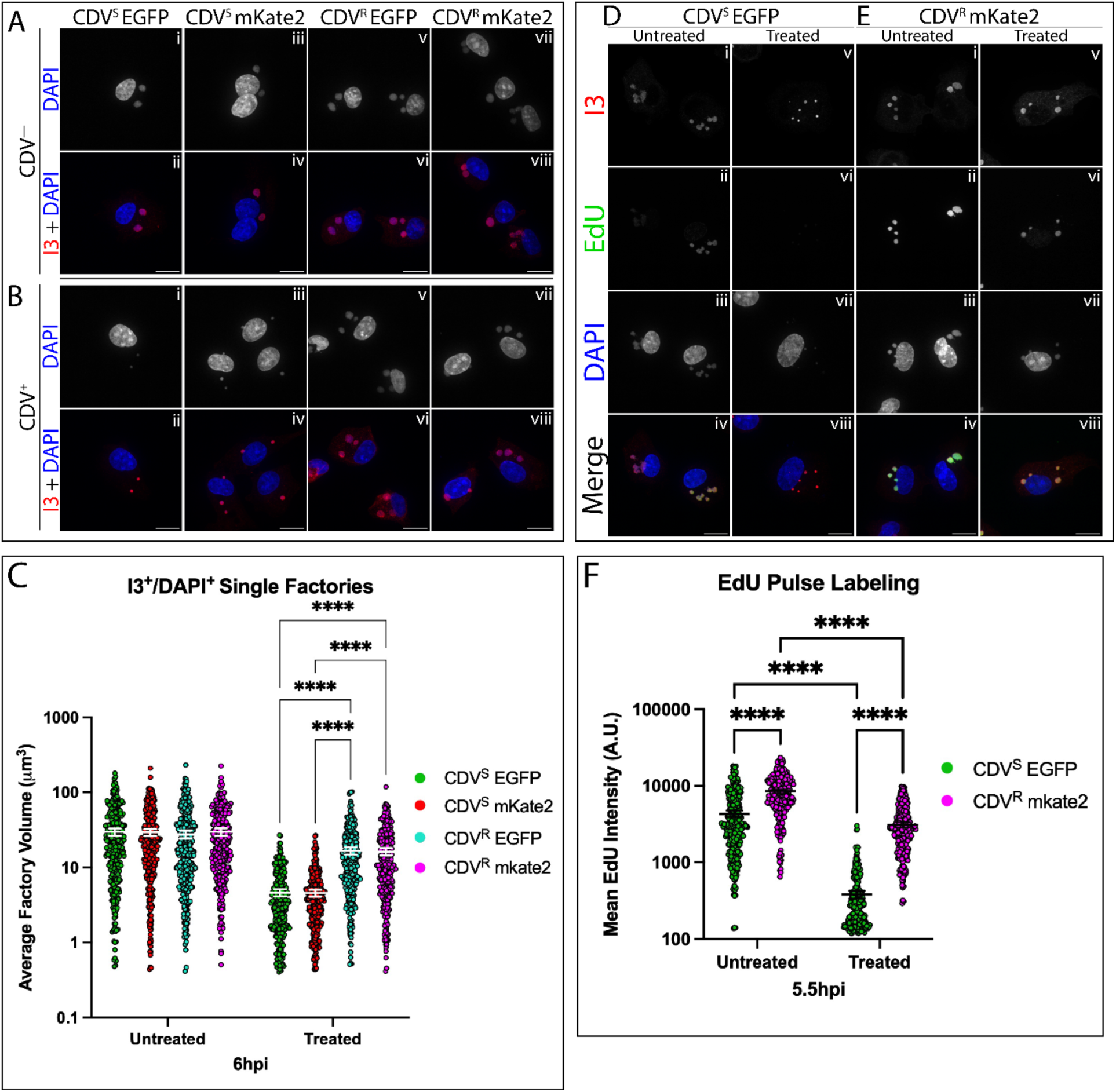
CDV inhibits DNA replication and prevents virus factory growth. A-B) BSC-40 cells were infected with the indicated VACV strains in the absence (A) or presence (B) of CDV. The cells were fixed at 6 hours post-infection and an antibody targeting the VACV I3 protein and DAPI dye were used to visualize virus factories. Cells were imaged by confocal microscopy. The z-stacks were merged based on fluorescence intensity and images were processed in Fiji. Scale bar = 15µm. C) Quantification from z-stacks of VACV factories in the absence or presence of CDV. The I3 antibody was used to identify virus factories in the cell, then the fluorescence intensity from DAPI was used to quantify the volume of the virus factory. The distributions include approximately 300 factory measurements collected across more than three independent biological experiments, and these data were analyzed using a two-way ANOVA with Tukey’s multiple comparisons. The mean and 95% confidence intervals are displayed. D-E) EdU pulse labelling of VACV-infected cells as a reporter for DNA replication. BSC-40 cells were infected with CDV^S^ EGFP (D) or CDV^R^ mKate2 (E). At 5 hours post-infection, the cells were pulsed with 10µm EdU. Labelling proceeded for 30 minutes before fixation. Copper-catalyzed click chemistry was used to conjugate Alexa Fluor™ 647 azide to EdU, then the cells were stained with an anti-I3 monoclonal antibody and DAPI dye. Cells were imaged by confocal microscopy. The z-stacks were merged based on fluorescence intensity and images were processed in Fiji. The EdU fluorescence was pseudo-coloured green. Scale bar = 15µm. F) Quantification of EdU incorporation into virus factories. The same protocol was used as in C except EdU fluorescence intensity was measured. The distributions include approximately 300 factory measurements collected across three independent biological experiments, and the data were analyzed using a two-way ANOVA with Šídák’s multiple comparisons. The mean and 95% confidence intervals are displayed.

To further demonstrate that CDV targets the DNA replication machinery, we used EdU pulse labeling and click chemistry to measure the rate of DNA synthesis in CDV^S^ and CDV^R^ factories (Figure 5D-F & Supplemental Figure 4). Pulsing the cells at 5 hours post-infection showed robust EdU incorporation into the virus factories after 30 minutes of labeling (Figure 5D, panel ii). However, there was almost no visible fluorescence from the EdU-AF647 molecule in the CDV^S^ factories when using CDV-treated cells (Figure 5D, panel vi). CDV^R^ virus factories still incorporated EdU in the presence of CDV, showing that E9 continued to synthesize DNA despite the drug treatment (Figure 5E panel ii & vi). Quantification of the fluorescent signal from EdU-AF647 provided concrete evidence that there was significantly more EdU incorporation into CDV^R^ factories than CDV^S^ factories under drug selection (Figure 5F). However, in untreated cells, we also observed that the CDV^R^ factories incorporated more EdU than the CDV^S^ factories. We speculated that the expression of different fluorescent proteins could have affected EdU incorporation, but switching the reporter viruses used in these experiments disproved this hypothesis (Supplemental Figure 5). Surprisingly, the CDV^R^ polymerase seemed more capable of accepting EdU as a substrate than the CDV^S^ polymerase, a finding that will be discussed below. Nevertheless, this experiment confirmed that CDV exerts an antiviral effect during DNA replication.

### The cro-EGFP and cro-mKate2 proteins do not uniquely label CDV^R^ or CDV^S^ virus factories

Having established a mechanism by which CDV inhibits factory growth, we proceeded to coinfection experiments using CDV^R^ and CDV^S^ viruses. In cells coinfected with CDV^R^ EGFP and CDV^S^ mKate2, we immediately noticed that cro-EGFP and cro-mKate2 proteins colocalized in the virus factories (Figure 6A-C). This was very surprising as previously, we found that in cells coinfected with viruses expressing EGFP-cro and mCherry-cro, the factories displayed uniform labeling with EGFP-cro or mCherry-cro, but not both proteins^31^. This posed a problem to using the cro-EGFP and cro-mKate2 proteins to track the CDV^S^ and CDV^R^ genomes as it was impossible to discern which factories originated from which virus. Nevertheless, we wanted to conduct live-cell imaging to see if there were any viruses in a coinfected cell that replicated significantly faster than the others, which might reflect one population of factories that originated from CDV^R^ genomes.

**Figure 6.**
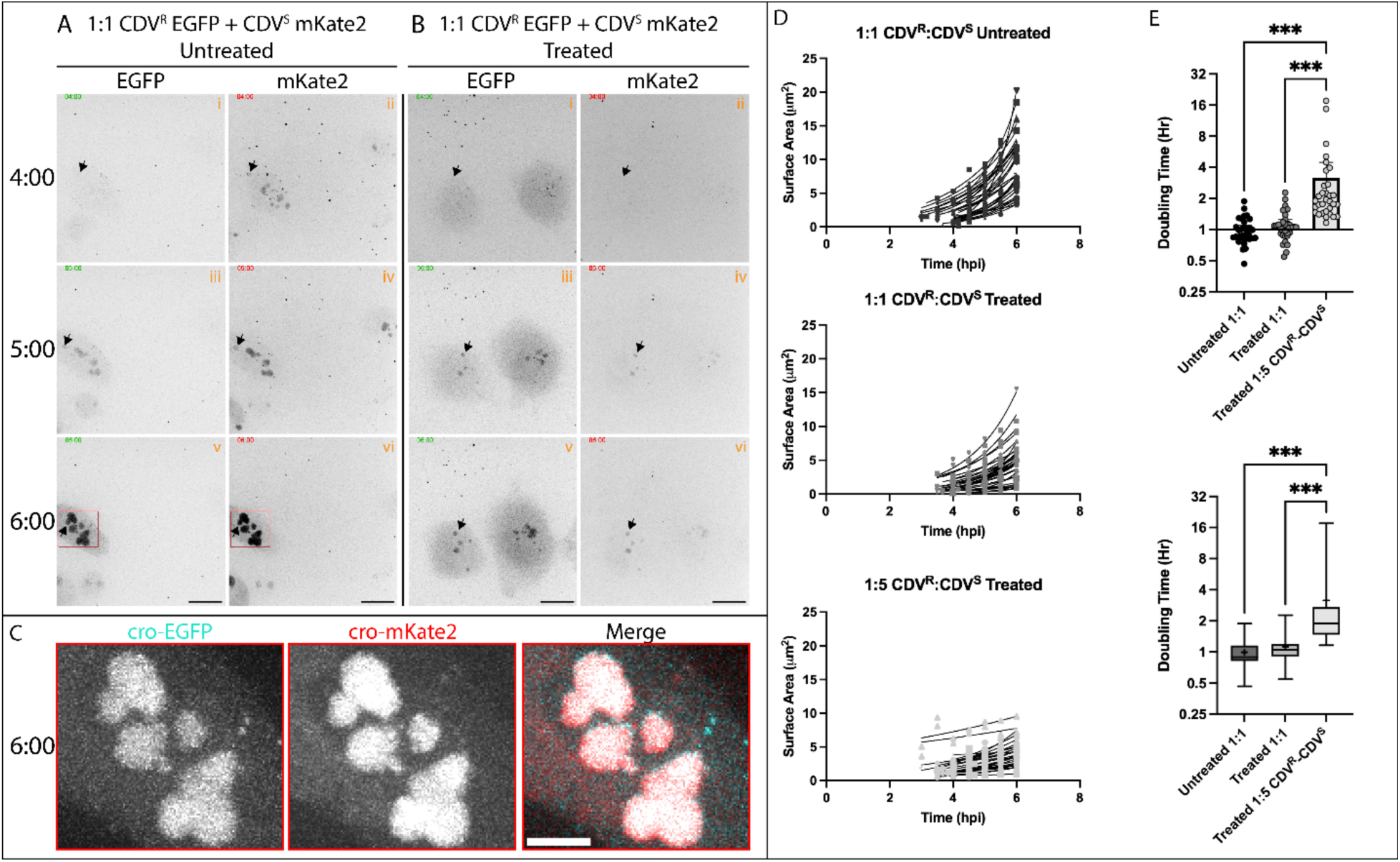
CDV^S^ and CDV^R^ viruses are indistinguishable by the expression of cro-FP proteins or by factory growth analysis . BSC-40 cells were synchronously coinfected at a combined MOI of 5 with VACV CDV^R^ EGFP and CDV^S^ mKate2 in the absence (A) or presence (B) of CDV. The infections were imaged from 3-6 hours post-infection. Representative images from more than three independent biological experiments are shown, and the EGFP and mKate2 fluorescence for each field of view are shown side-by-side. Arrows indicate factories that did not fuse with adjacent factories throughout the experiment. Scale bar = 15µm. C) The insets from panels v and vi of part A and a merged image are shown. The fluorescent signals from cro-EGFP and cro-mKate2 inside the virus factories overlap, preventing identification of CDV^R^ and CDV^S^ viruses. Scale bar = 5µm. D) Quantification of live-cell imaging of virus factories from cells coinfected with VACV CDV^S^ EGFP or CDV^R^ mKate2. >30 virus factories were measured from each coinfection model across >3 independent biological experiments. E) The doubling times of the non-linear regression curves shown in D were compiled and analyzed using a one-way ANOVA with Tukey’s multiple comparisons. The data were plotted as a box-and-whisker plot in E to show the normal distribution of the data (whiskers). + = mean, ̶ = median, box = first and third quartile, whisker = minimum and maximum.

### The growth rate of the virus factories in coinfected cells is dependent on the relative gene doses of CDV^R^ and CDV^S^ polymerases

In the absence of CDV, the factories in cells coinfected with CDV^R^ EGFP and CDV^S^ mKate2 rapidly grew during the time course (Figure 6A). We observed stronger fluorescence intensity in the mKate2 channel, indicating that there could have been more CDV^S^ viruses infecting the cell. In CDV-treated cells, all the factories seemed to be growing at the same rate (Figure 6B). After CDV treatment, there was no obvious change in the growth kinetics of the virus factories compared to the untreated cells (Figure 6D). To accentuate any differences in growth kinetics between CDV^S^ and CDV^R^ viruses, we conducted a coinfection experiment in which there were five CDV^S^ viruses to one CDV^R^ virus entering the cell (Supplemental Figure 7). We hypothesized that if the CDV^R^ virus maintained its competitive advantage over the CDV^S^ virus, then we would see ⅙ of the virus factories in a distribution growing significantly faster than the rest. However, we noticed a flattening of the growth curves when there were excess CDV^S^ viruses in the cell (Figure 6D). Extracting and analyzing the doubling times of the factories showed that when the CDV^R^ and CDV^S^ viruses are added to the cells at a 1:1 ratio, there was no difference in the growth of the factories between untreated and treated cells (Figure 6E). When there were excess CDV^S^ viruses, the mean doubling time was significantly higher than when the viruses were at a 1:1 ratio (p<0.001). We also plotted the data as a box-and-whisker plot and found that there were no factories outside of the normal distribution of the data for any of the infection conditions (Figure 6E).

### CDV^S^ viruses parasitize the growth advantage of CDV^R^ viruses in coinfected cells

Collectively, these observations suggested that CDV^R^ and CDV^S^ viruses replicated at the same rate in coinfected cells, causing the CDV^R^ virus to lose the competitive advantage it possessed over CDV^S^ viruses in single infections. However, the colocalization of cro-EGFP and cro-mKate2 prevented us from directly measuring the growth of CDV^S^ and CDV^R^ viruses. Therefore, we developed a probe for fluorescence *in situ* hybridization (FISH) that could differentiate CDV^S^ and CDV^R^ viruses in a coinfected cell (Figure 7A). The probe was complementary to the cro-EGFP gene. In this experiment, we used CDV^S^ EGFP and VACV VDG1.3, which was the parental strain used to make CDV^R^ EGFP and CDV^R^ mKate2. Because VDG1.3 did not have any insertions into *J2R*, the probe target sequence was only inside the CDV^S^ factories. In coinfected cells, this resulted in the probe hybridizing to only a portion of the factories in the infected cells (Figure 7B, panel vi & xiv). Additionally, we could not detect any strong hybridization inside the nuclei, which showed that either the probe did not hybridize to DNA without the target sequence, or the hybridization and washing protocol successfully eliminated non-specific hybridization (Supplemental Figure 8). Based on the hybridization pattern, we assigned the factories as either CDV^S^ or CDV^R^ (Supplemental Figure 8) and quantified the size, showing that in CDV-treated cells, there was no significant difference between the size of the CDV^S^ and CDV^R^ virus factories (Figure 7C). This experiment confirmed that in coinfected cells, the CDV^R^ and CDV^S^ viruses replicated at the same rate, explaining why in our previous experiments, the CDV^S^ virus survived the selective pressure in high-MOI coinfections, but not when the two strains replicated in isolation (Figure 2).

**Figure 7.**
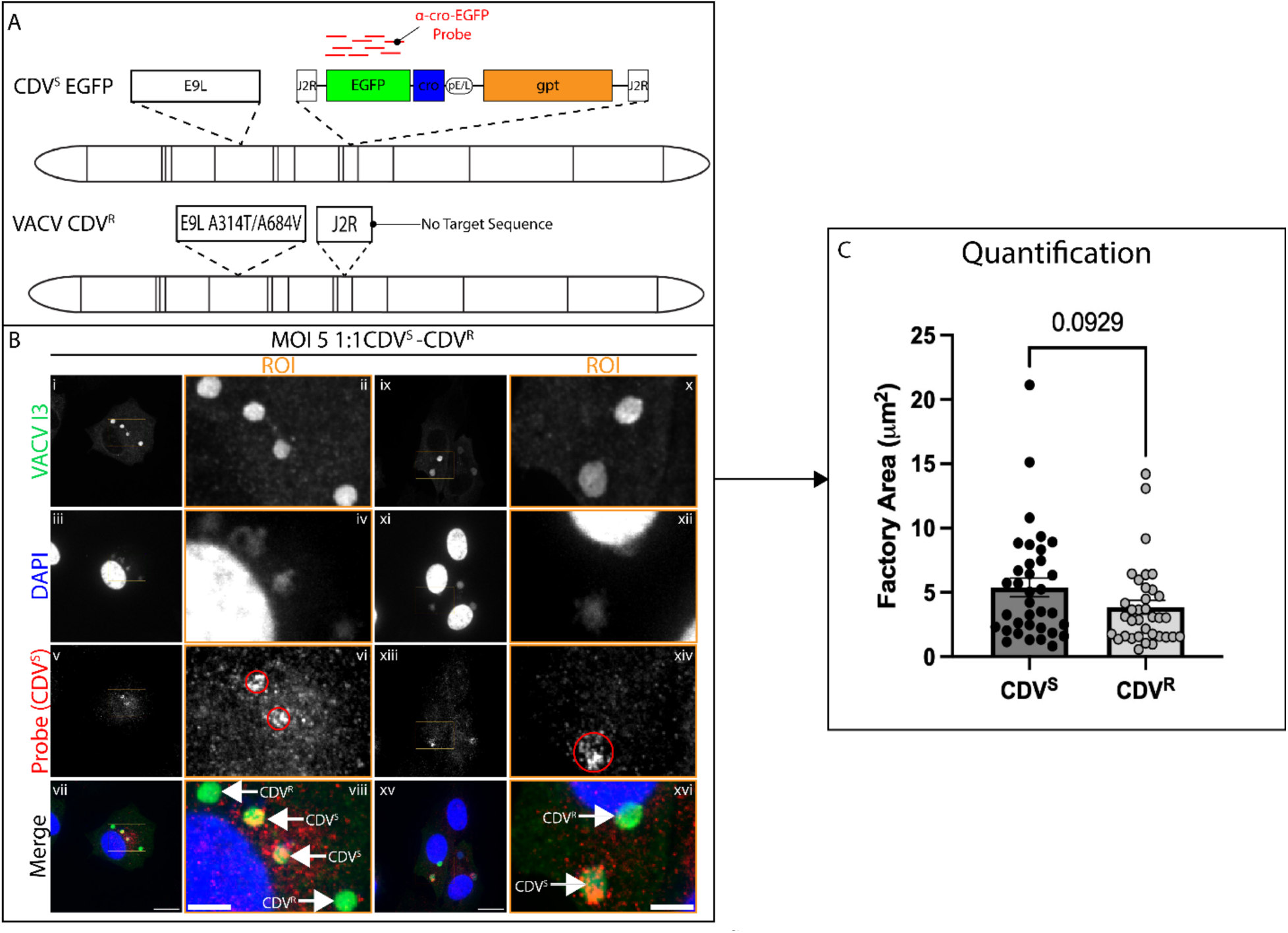
FISH imaging shows that CDV^R^ and CDV^S^ viruses replicate at the same rate in coinfected cells. A) A DNA FISH probe was generated that specifically targeted the cro-EGFP sequence in the CDV^S^ EGFP genome (see Materials and Methods). Coinfection with the parental CDV^R^ virus that did not express cro-EGFP allowed specific labeling of the CDV^S^ virus in coinfected cells. B) BSC-40 cells were treated with 330µM CDV and synchronously coinfected with VACV VDG1.3 (CDV^R^, no fluorescent protein expression) and VACV CDV^S^ EGFP. Cells were fixed at 6 hours post-infection in 4% PFA overnight before in-situ hybridization. After hybridization, IF was used to visualize the I3 protein and DAPI counterstaining was added. Two representative images are shown with the inset showing a region of interest (ROI) beside the full image. The panels show virus factories where the probe hybridized to the genomic DNA (red circles), showing that the virus factories contain the CDV^S^ genome. Adjacent virus factories where the probe did not strongly hybridize likely originated from the coinfecting CDV^R^ virus. Scale bar = 15µm, ROI scale bar = 5µm. C) CDV^S^ and CDV^R^ viruses identified by FISH were quantified. Factories were measured in 2D as for live-cell imaging using a maximum intensity projection of a z-stack. The distributions contain >30 virus factories imaged from two independent biological experiments, and an unpaired, two-way t-test was used for statistical analysis.

## Discussion

A generally accepted paradigm is that DNA viruses evolve at a slower rate than RNA viruses. However, there is a growing body of work showing that poxvirus evolution is rapid and dynamic, owing to both high rates of mutation and diverse molecular strategies to adapt to selective pressures^2^. Our data showed that a trait like CDV^R^ can undergo rapid selection after a single round of replication when the viruses in the population enter separate cells (Figure 2). However, the presence of coinfecting CDV^S^ viruses interfered with this selection and prevented the rapid outgrowth of CDV^R^ mutants (Figure 2). In terms of population fitness, under selection from CDV, a scenario in which all viruses replicate in isolation promoted fitness, whereas a coinfection negated this purifying selection. Thus, it appears that at least for CDV^R^, coinfection between CDV^R^ and CDV^S^ viruses is detrimental to the population, as the coinfection will maintain the less fit CDV^S^ strain in the population.

When CDV^S^ and CDV^R^ viruses replicated in isolation, there was a clear difference in the kinetics of virus factory growth (Figures 3 and 4). The CDV^S^ virus factories displayed significantly slower growth under drug selection compared to CDV^R^ virus factories (Figure 4). EdU pulse labeling showed that the CDV^S^ polymerase stopped synthesizing DNA after CDV treatment, whereas the CDV^R^ polymerase resisted this effect (Figure 5). This experiment also showed that the CDV^R^ polymerase incorporated more EdU into the replicated DNA than the CDV^S^ polymerase (Figure 5F, Supplementary Figure 5). The A684V substitution in the nucleotide binding pocket may have caused this phenotype, as this substitution induces a mutator phenotype^12,35^. Position 684 may contribute to positioning residues that bind to incoming nucleotides^16,17^; while this disfavours CDV entering the active site, it is possible that the A684V substitution could enhance EdU entrance into the active site.

The loss of DNA synthesis ultimately led to smaller virus factories in CDV-treated cells infected with the CDV^S^ virus compared to cells infected with the CDV^R^ virus (Figure 5). These data showed that CDV exerts an antiviral effect during DNA replication, consistent with the data from primer extension assays using purified VACV DNA polymerase holoenzyme complexes^7,8^. However, this does not rule out additional mechanisms of action downstream from DNA replication. Atomic force microscopy showed that the structure of viral DNA is severely distorted after CDV treatment^33^. Previous work using nuclear magnetic resonance showed that CDV molecules disrupt the DNA duplex, owing to the acyclic structure of CDV^36^. A structural defect from incorporating CDV into the daughter strand would likely have far-reaching effects on gene expression, protein synthesis, and assembly.

In coinfected cells, the CDV^S^ virus parasitized the growth advantage of the CDV^R^ virus in CDV-treated cells (Figures 6 and 7). This suggested that the CDV^S^ factories somehow acquired the functional CDV^R^ polymerase from the coinfecting virus. The colocalization of cro-EGFP and cro-mKate2 further showed that proteins encoded by one virus are not strictly retained in the factory of genetic origin (Figure 6C). Several previous observations suggest that this phenotype is expected to arise at early stages of replication. After entry, the capsid is transcriptionally active and extrudes early transcripts into the cytoplasm^37–39^. The transcripts localize to polysome-dense aggregates that are distinct from the uncoated viral genome^40,41^. This means that early protein synthesis occurs in the cytoplasm, not in the virus factory. DNA-binding proteins like the DNA polymerase and associated factors likely return to the genome through electrostatic interactions driving their diffusion. Thus, there is nothing that can constrain a CDV^R^ polymerase from binding a CDV^S^ genome, and vice versa (Figure 8A). Tagging the individual E9 proteins with epitope tags and conducting immunofluorescence microscopy would be one way to obtain concrete evidence for this model. Combining this approach with RNA FISH could provide greater insight into the pre-factory stage of the poxvirus lifecycle.

**Figure 8.**
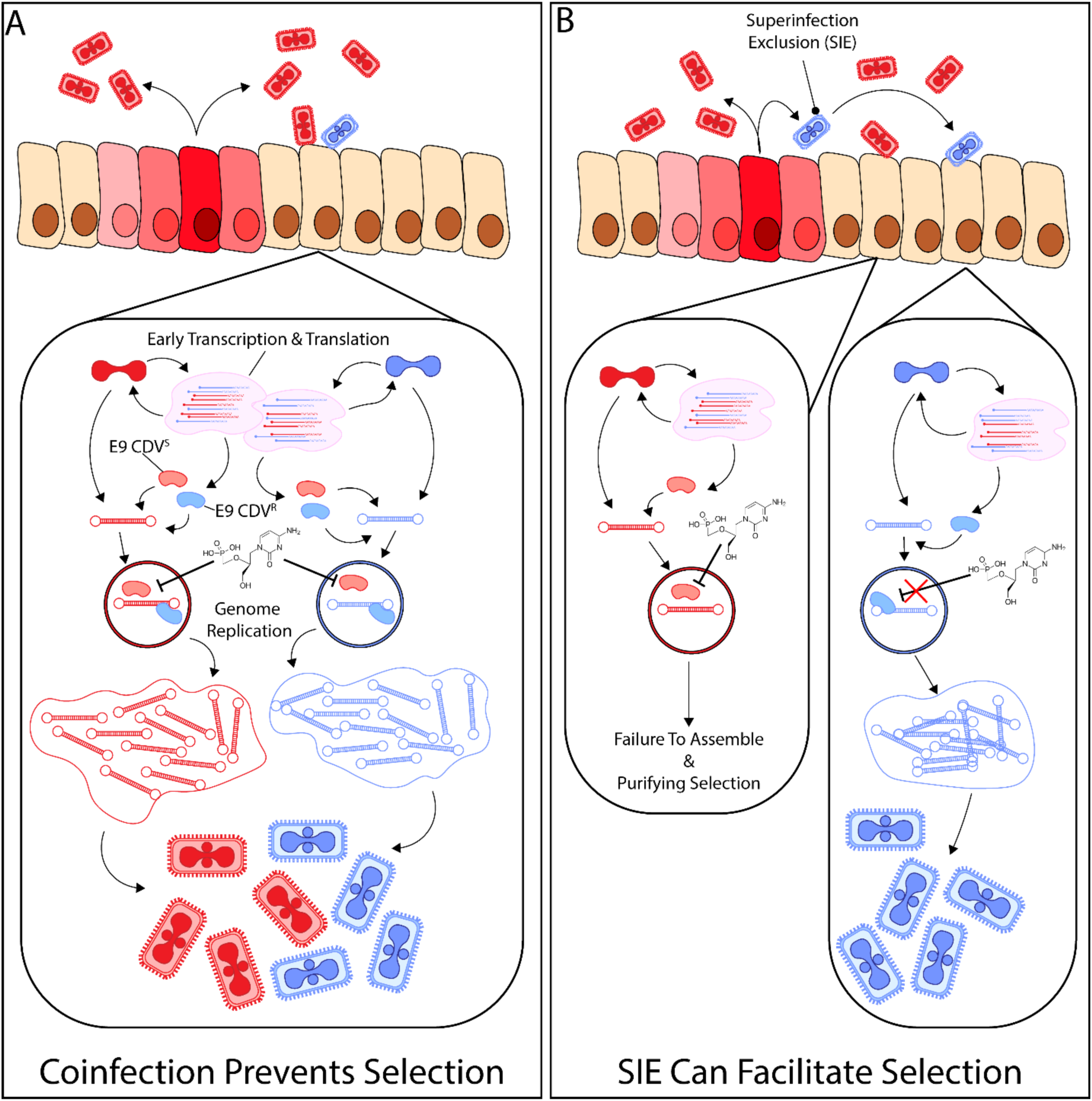
Model for the mechanism of selection for CDV^R^. A) When CDV^S^ (Red) and CDV^R^ (Blue) viruses infect the same cell, the two opposing traits become codominant with each other. During uncoating, the CDV^S^ and CDV^R^ polymerases are synthesized in the cytoplasm. During factory formation, the genomes randomly sequester both forms of the polymerase. The presence of CDV^R^ polymerases in the virus factories is sufficient to replicate enough copies of the genome to proceed to late stages of infection. Equal numbers of CDV^S^ and CDV^R^ progeny assemble. B) When the CDV^S^ and CDV^R^ viruses infect separate cells, the CDV^S^ virus cannot parasitize the growth advantage of the CDV^R^ virus. This allows the CDV^R^ virus to outcompete the CDV^S^ virus and purifying selection to operate in the population. One possibility is that a mechanism such as superinfection exclusion prevents interference from coinfections, although this phenomenon requires more investigation.

In our experiments, we artificially forced coinfection by using an MOI of 5 or 10. However, it is unclear how often coinfection with multiple poxviruses occurs *in vivo*. The circulation of recombinant strains in populations of different poxviruses suggests that coinfection can occur *in vivo*^42–47^. *In vitro*, there is evidence that poxviruses actively prevent superinfection, which would naturally reduce the probability of coinfection^48^. Two early VACV proteins, A33 and A36, work to capture incoming particles and transfer them to actin filaments that push them towards uninfected cells^49^. The authors described this phenomenon as a mechanism that enhances the spread of the virus in a tissue. However, this could also act as a mechanism to maintain population fitness (Figure 8B). Superinfection exclusion could prevent the entry of defective particles into cells infected with replication-competent virus, thus preventing the accumulation of non-infectious particles. Laliberte and Moss also described SIE in this context, albeit through a distinct mechanism^50^. Unfortunately, this model has limitations, primarily that we do not understand why poxviruses evolved to prevent superinfection. To our knowledge, the studies showing SIE have only investigated this phenomenon *in vitro*. These experiments have the luxury of artificially increasing the MOI and changing the timing of the second virus compared to the first infection. These models are invaluable tools for mechanistic work, but they do not recapitulate the dynamics of a virus infecting and spreading in a complex host tissue. Further work is needed to investigate this phenomenon and its evolutionary benefits to poxvirus populations. Nevertheless, our data do show that coinfection has a negative effect on population fitness, allowing less-fit viruses to survive in purifying selection. Brincidofovir (BDV) and CDV are important therapeutics for treating orthopoxvirus infections in patients^4^. The alternative therapeutic, tecovirimat, has not shown superior efficacy in reducing symptom duration compared to supportive care in recent clinical trials^51–53^. It appears moderately effective in treating infection, but there is a very low barrier to resistance^10,11,54^. *In vitro*, a single passage in the presence of tecovirimat can select for highly resistant viruses^55^. In another study, the selection took more than 14 passages, though it is difficult to properly compare the two studies with different methodologies^5^. Generally, it is thought that the mutational barrier to CDV^R^ is high, as initial attempts to isolate resistant viruses required extensive passage^12^. However, our data showed that CDV^R^ viruses rapidly outcompete CDV^S^ viruses in the population when the two strains replicate in isolated cells (Figure 2). These data suggest that as a virus population continues circulating, there can be rapid selection for novel, advantageous traits like drug resistance within a single round of viral replication. This is consistent with work studying adaptations during spillover, wherein the poxvirus genome becomes remarkably plastic, duplicating protein kinase R antagonists to increase viral fitness within a few passages^2,56–58^. For CDV^R^, SIE could represent an additional molecular mechanism for rapid evolution as well as maintaining fitness as the virus continuously circulates.

As MPXV continues to spread in humans, there are more chances for adaptive mutations to become fixed in the population of circulating viruses. The continued re-emergence of MPXV is also increasing the use of BDV and CDV, creating new selective pressures and potentially selecting for resistance mutations. While the specific mutations we used attenuated the virus *in vivo*, there could possibly be compensatory mutations that improve fitness. The rapid outgrowth of such mutants would pose a significant threat to containment efforts aimed at curbing the spread of MPXV through antiviral drug use. Ultimately, our data suggest there is a greater need for additional antivirals and combination therapies that will reduce the risk of developing drug resistance. Such efforts would support pandemic preparedness as poxviruses continue to spillover and adapt to new selective pressures.

## Materials & Methods

### Cells and Viruses

African green monkey kidney epithelial cells (BSC-40) were purchased from the American type culture collection (ATCC) and grown in minimal essential medium (MEM) supplemented with non-essential amino acids, L-glutamine, sodium pyruvate, antibiotic/antimycotic, and 5% FetalGro, all of which were purchased from Thermo Fisher Scientific. Our lab stock of VACV Western Reserve (WR) was originally purchased from ATCC and a clonal population was previously generated by plaque purification. VACV VDG1.3 was generated as previously described and is a non-clonal WR strain with the CDV^R^ mutations in the *E9L* gene^12^. VACV titers were determined using a plaque assay on BSC-40 cells.

### Recombinant Virus Construction

To generate recombinant viruses expressing cro-EGFP or cro-mKate2, the synthetic genes were placed under the control of a consensus poxvirus early/late promoter (pE/L) and cloned into the pTM3 vector, a gift from Dr. Bernard Moss with homology to the *J2R* gene of vaccinia and an *Escherichia coli* xanthine-guanine phosphoribosyltransferase cassette (gpt). BSC-40 cells were infected with VACV WR or VACV VDG1.3 at an MOI of 0.5. 2 hours post-infection, the cells were transfected with 2ug of linearized pTM3-pE/L-cro-EGFP or pTM3-pE/L-cro-mKate2. 2 days post-infection, the monolayer was harvested and subjected to three freeze/thaw cycles to release infectious virus. Recombinant viruses (EGFP^+^ or mKate2^+^, gpt^+^) were plaque purified by infecting fresh BSC-40 cells and overlaying the cells with MEM supplemented with 1.7% Noble agar, and mycophenolic acid (MPA) selection media. Purified recombinant virus stocks were amplified in BSC-40 cells and purified by centrifugation through a 36% sucrose cushion as previously described^59^.

### Multi-Step Growth Curves

BSC-40 cells were seeded in 6-well plates and infected with different VACV strains at an MOI of 0.03. The monolayers were harvested at 0, 3, 6, 12, 24, and 48 hours post-infection. Infectious virus was released by freeze/thaw cycles and titrated on BSC-40 cells.

### Plaque Reduction Assays

BSC-40 cells were seeded in 12-well plates and infected with roughly 100 plaque-forming units (PFU) of different VACV strains. One-hour post-infection, the inoculum was removed and MEM with 1% w/w carboxymethylcellulose and increasing concentrations of CDV was added. Forty-eight hours post-infection, the monolayers were fixed and stained with crystal violet stain (1.3% w/v crystal violet, 5% v/v EtOH, 11% v/v formaldehyde). Plaques were counted and normalized to untreated wells to calculate the plating efficiency.

### Fluorescence Microscopy

For basic imaging of virus plaques, infected cells were imaged using an EVOS FL Auto Imaging System (Thermo Fisher) using FITC or TRITC lasers and a 10X objective. For imaging whole well plates, a BioTek Cytation was used with FITC and TRITC lasers at 1.25X objective. Autoexposure was used to determine the optimal exposure settings, which were then used for all subsequent imaging experiments.

All live-cell imaging was conducted using a WaveFx spinning disc confocal microscope mounted on an Olympus IX-81 microscope base and a Hamamatsu C9100-13 EMCCD camera. For live-cell experiments, BSC-40 cells were seeded into 35mm glass-bottom dishes in FluoroBrite^TM^ media supplemented with 2% fetal bovine serum and non-essential amino acids. For experiments with CDV treatment, CDV was thawed and diluted to 330uM in the media at the time of seeding the cells. The cells were synchronized on ice for 15 minutes before inoculation with different VACV strains at an MOI of 5. The viruses were allowed to bind to the cells for 1 hour at 4°C before the inoculum was replaced with warmed 1x MEM. The cells were replaced to 37°C with 5% CO_2_ before mounting in a heated live-cell chamber. Live cells were imaged with a 60X/1.42 oil lens. Fluorophores were excited with either a 491nm or 561nm single wavelength laser.

For fixed samples, BSC-40 cells were seeded on 15mm glass coverslips in 1x MEM supplemented with 2% FBS. For experiments with CDV treatment, CDV was thawed and diluted to 330uM in the media at the time of seeding the cells. The cells were synchronously infected with different VACV strains. At different timepoints, the infected cells were washed in 1X PBS before fixation in 2% paraformaldehyde for 10 minutes at room temperature. Free aldehydes were quenched in 0.1M glycine in PBS pH 7.4 for 20 minutes. Cells were permeabilized in 0.2% TritonX-100 for 2 minutes at room temperature, then washed in 1X PBS three times before blocking with 3% bovine serum albumin in 1x PBS pH 7.4 for 1 hour at room temperature. Primary antibody staining was done for 2 hours at room temperature. The cells were then washed in 1X PBS-T three times before adding the secondary antibody and DAPI (10µg/mL) stain for 1 hour at room temperature. The cells were washed in 1X PBS-T and mounted in SlowFade mounting medium (Thermo Fisher).

For experiments with click chemistry, the cells were pulsed with 10µM EdU at 5 hours post-infection. After 30 minutes, the cells were fixed and permeabilized. The click chemistry reaction (86mM Tris-HCl pH 9, 4mM CuSO_4_, 4.8µM AlexaFluor^TM^ 647 Azide, 10mM sodium-L-ascorbate) was prepared fresh and added to the cells for 30 minutes before washing the cells in 1x PBS-T and proceeding with the same staining protocol described above.

#### Fluorescence *in situ* hybridization

FISH probes were prepared from the plasmid pMK-QR-cro-EGFP by nick translation using an ARES® 647 kit (Molecular Probes), which incorporates a 5-(3-aminoallyl)-dUTP molecule into the products. The reactions were assembled following the manufacturer’s instructions and successful nick translation was assessed by running the products on a 0.8% agarose gel in 1x Tris acetate EDTA buffer. The resulting DNA fragments were purified using a QIAquick PCR Purification Kit (QIAGEN) and labeled with AlexaFluor^TM^ 647 succinamide carboxylic acid. The labeled probes were purified again, and the purity and concentration were assessed using a NanoDrop spectrophotometer.

For FISH experiments, BSC-40 cells were seeded on 18mm glass coverslips in 1X MEM with 2% FBS and 330uM CDV. Cells were synchronously coinfected with VACV VDG1.3 (CDV^R^, no cro-EGFP insert) and VACV pE/L-cro-EGFP (CDV^S^ EGFP, has the FISH target sequence in *J2R*) at a combined MOI of 5. At 6 hours post-infection, the cells were fixed in 4% PFA overnight at 4oC. Free aldehydes were quenched and the cells were permeabilized before treatment with RNase. The cells were then blocked and the FISH probe was diluted in hybridization buffer (50% v/v formamide, 5% v/v dextran sulfate, 0.2x saline sodium citrate pH 7, 50mM sodium phosphate pH 7, 1mM EDTA, 0.01mg/mL salmon sperm DNA, 1x Denhardt’s Reagent, 2ng/µL probe). After blocking, the coverslips were placed into SecureSeal^TM^ Hybridization chambers and roughly 100uL of hybridization buffer was added. Pre-hybridization was done at 70°C for 5 minutes, then the cells were snap-cooled in an ice water bath and placed in a 50°C incubator overnight. Following hybridization, the cells were washed twice in 2X SSC with 50% formamide for 30 minutes at 50°C, then washed with PBS extensively before immunofluorescence staining to compare the localization of the FISH probe with VACV I3 protein and DAPI-stained viral DNA.

### Image Analysis

For analyzing live-cell imaging data, the movies were extracted from Volocity v6.3 as tagged image file (TIF) stacks and analyzed in FIJI^60^as previously described^30^. FISH images were also analyzed in FIJI.

For measuring virus factory volume, the “compartmentalize” tool from Volocity was used as previously described^61^. Briefly, virus factories were identified based on the fluorescent signal from the primary/secondary antibody combination targeting the I3 protein. Virus factories were assigned as “containers” and subsequently, DAPI-stained “objects” were identified within the virus factory. This process eliminated nuclei from any measurements. Automatic thresholding of the fluorescence values was used for each image to minimize external manipulation of the analysis. This protocol was applied to all images taken and then checked to ensure that the software did not make any mistakes (e.g. two factories that were touching could be identified as a single factory by the protocol). The volume measurements were then extracted into GraphPad Prism. For measuring EdU fluorescence intensity, the same protocol was used except that instead of identifying the “objects” using DAPI fluorescence, the fluorescent signal from EdU was used.

For measuring fluorescence intensity as a line profile, the “line profile” tool from Volocity was used and fluorescence values were extracted. The mean fluorescence intensities of the three channels were corrected for background signal using the black limit of the entire image. A background correction was also applied to the white limit to reflect the total range of values within each channel. All values were then normalized to the corrected white limit to generate a percentage of the total fluorescence within the line. This was done so that all three channels could be plotted within a single graph and to account for differences in the strength of fluorescence between the three channels that might cause scaling problems when graphing. The data were then plotted in GraphPad Prism.

### Statistics

All statistical analyses were performed in GraphPad Prism version 9.3.1. The threshold for significance was p<0.05. Significance was represented in figures as follows: * p<0.05, ** p<0.01, **** p<0.001. All analyses used three independent biological replicates except for the FISH experiment, which included two biological replicates. Imaging analysis was done by pooling measurements compiled from at least three biological replicates. This was done to capture the natural variation in a monolayer of infected cells that would be lost if individual factories were treated as technical replicates. At least 30 factory measurements were included for live-cell imaging experiments and for the FISH experiment. The only exception to this is in Figure 4. Two factories were removed from the “CDV^S^ Treated” group because Prism generated growth curves with negative values for the doubling time. IF experiments included approximately 300 factories.

## Acknowledgements

Microscopy experiments were performed at the University of Alberta Faculty of Medicine & Dentistry Cell Imaging Core, which receives financial support from the Faculty of Medicine & Dentistry and Canada Foundation for Innovation (CFI) awards to contributing investigators.

This work was supported by grants from the Canadian Institutes of Health Research (to D.H.E and R.S.N), The Natural Sciences and Engineering Research Council (to D.H.E), the Li Ka Shing Institute of Virology (to R.S.N and D.H.E), and the Alberta Ministry of Technology and Innovation through SPP-ARC (Striving for Pandemic Preparedness – The Alberta Research Consortium (to R.S.N). Studentship support was received from the Li Ka Shing Institute of Virology and the Faculty of Medicine and Dentistry (to S.Z.L).

## Author Contributions

R.S.N, D.H.E, and S.Z.L conducted conceptualization. R.S.N and D.H.E conducted funding acquisition and project administration. S.Z.L designed the methodology. S.Z.L and Y-C.L conducted the investigation, data curation and formal analysis for the experiments. S.Z.L wrote the original draft with significant contributions from R.S.N and D.H.E, and Y-C.L, D.H.E and R.S.N reviewed and edited the manuscript.

## Notes

### Competing Interest Statement

The authors have declared no competing interest.

