## Supplemental Figures for "The selection for cidofovir-resistant mutant vaccinia viruses is inhibited by coinfecting cidofovir-sensitive wildtypes"

**Supplemental Figure 1. Insertion of cro-mKate2 or cro-EGFP into the J2R does not reduce virus fitness but changes CDV sensitivity**

**Supplemental Figure 2. Titration of progeny from low- and high-MOI competition assays**

**Supplemental Figure 3. Additional IF images of CDV$^{S}$ and CDV$^{R}$ factories**

**Supplemental Figure 4. Additional IF images of EdU-labeled CDV$^{S}$ and CDV$^{R}$ virus factories**

**Supplemental Figure 5. CDV$^{R}$ virus factories show higher levels of EdU incorporation than CDV$^{S}$ virus factories**

**Supplemental Figure 6. Live-cell imaging of cells coinfected with CDV$^{R}$ EGFP and CDV$^{S}$ mKate2**

**Supplemental Figure 7. Live-cell imaging of coinfected cells**

**Supplemental Figure 8. Line profiles of VACV infected cells used for FISH**

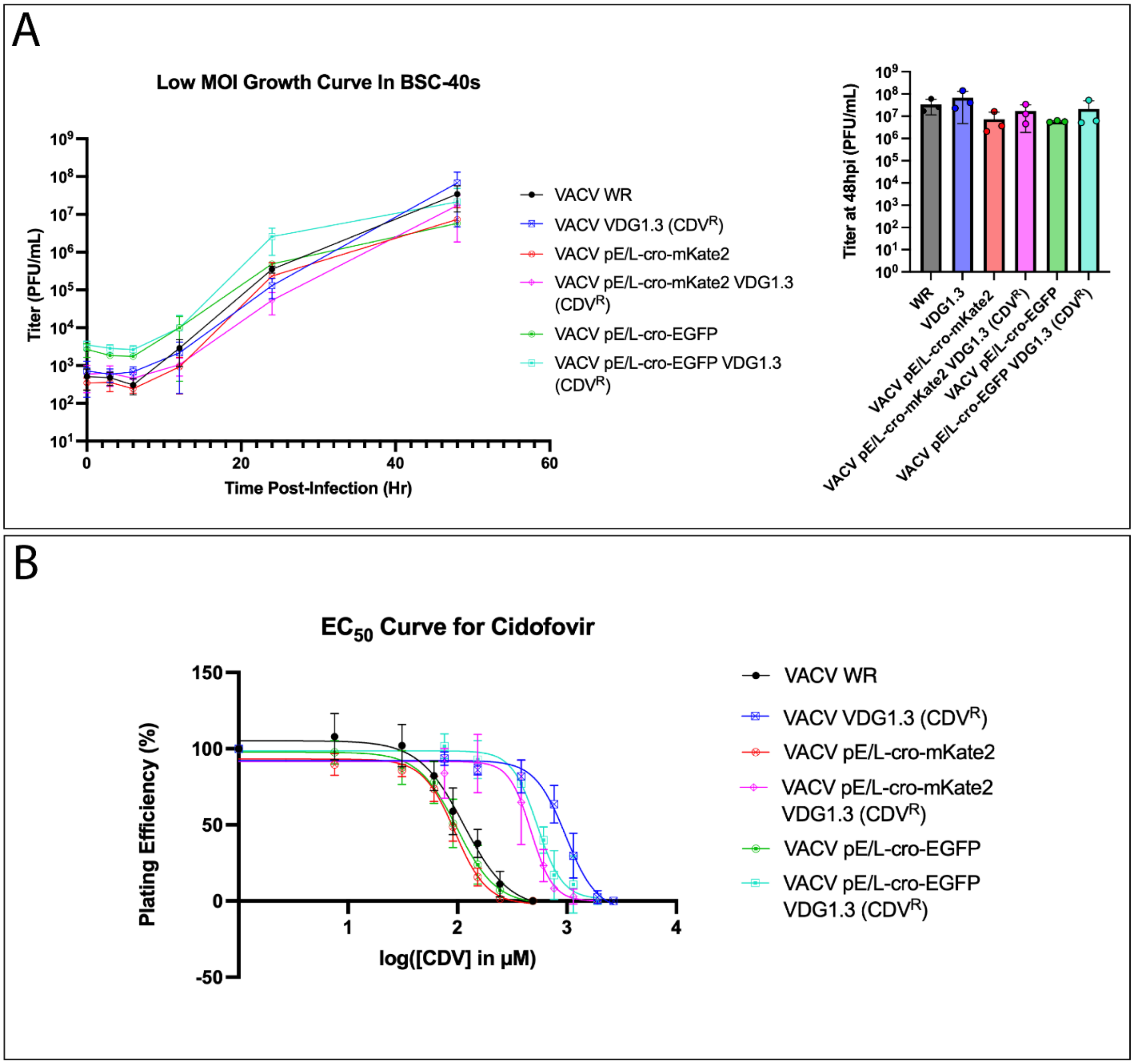


**Supplemental Figure 1. Insertion of cro-mKate2 or cro-EGFP into the J2R does not reduce virus fitness but changes CDV sensitivity.** A) BSC-40 cells were infected with the indicated viruses at an MOI of 0.01. The cell monolayers were harvested at the indicated timepoints, and the progeny were released from the cells by repeated freeze/thaw cycles. The progeny were titrated on fresh BSC-40 cells. For quantification, the titers of each virus at 48 hours post-infection were analyzed using a one-way ANOVA with Tukey's multiple comparisons. The mean ± standard deviation from three independent biological experiments is displayed. B) Roughly 100 PFU of the indicated viruses were plated on BSC-40 cells with increasing concentrations of CDV. 48 hours post-infection, the plaques were counted and normalized to untreated wells to calculate plating efficiency. Data were plotted in GraphPad Prism and non-linear regression was used to generate a best-fit curve to the data. The mean ± standard deviation from three independent biological experiments is displayed.

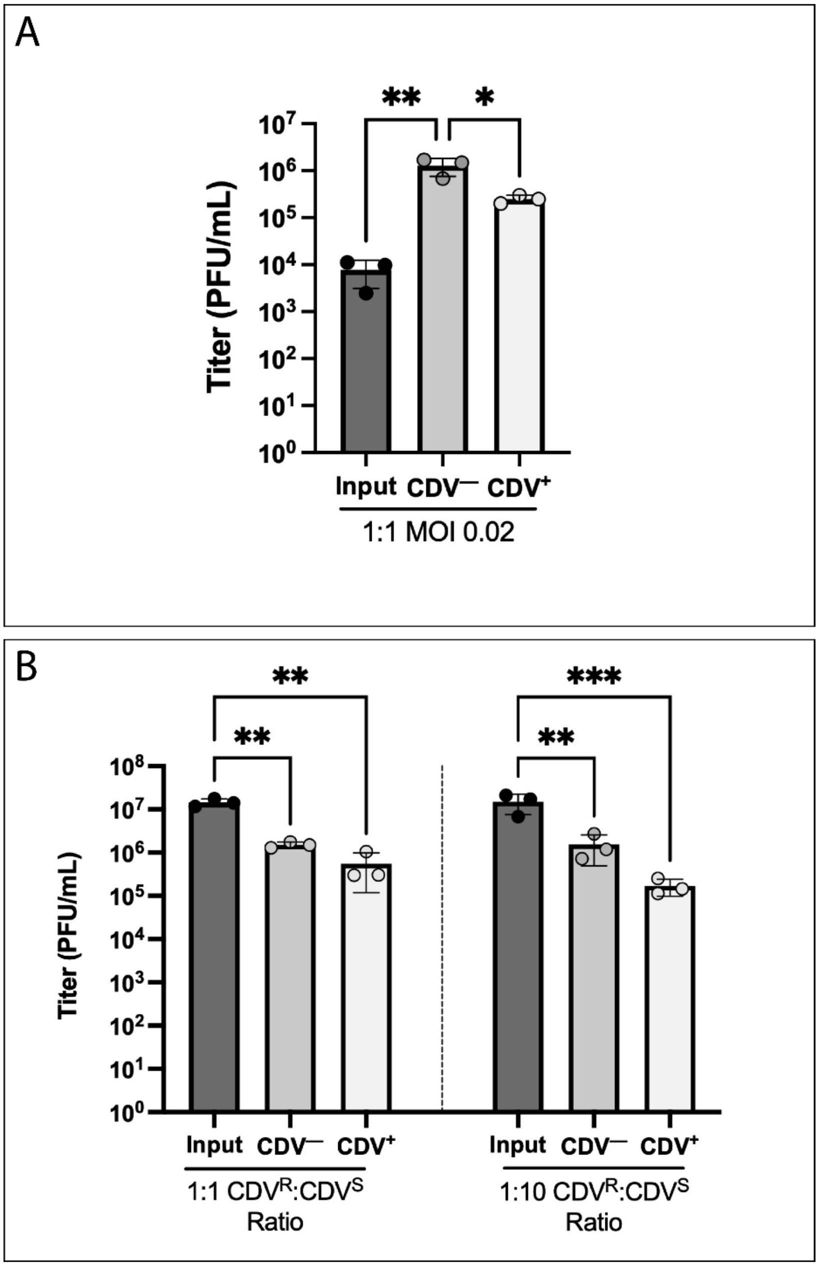


**Supplemental Figure 2. Titration of progeny from low- and high-MOI competition assays.** The input viruses and progeny from cells infected with VACV $CDV^R$ EGFP and $CDV^S$ mKate2 were titrated on BSC-40 cells, and the data analyzed using a one-way ANOVA with Tukey's multiple comparisons. A) A low MOI (0.02) was used to model a scenario in which the two viruses replicate in isolation. B) A high MOI (10 or 11) was used to model a coinfection between the two viruses.

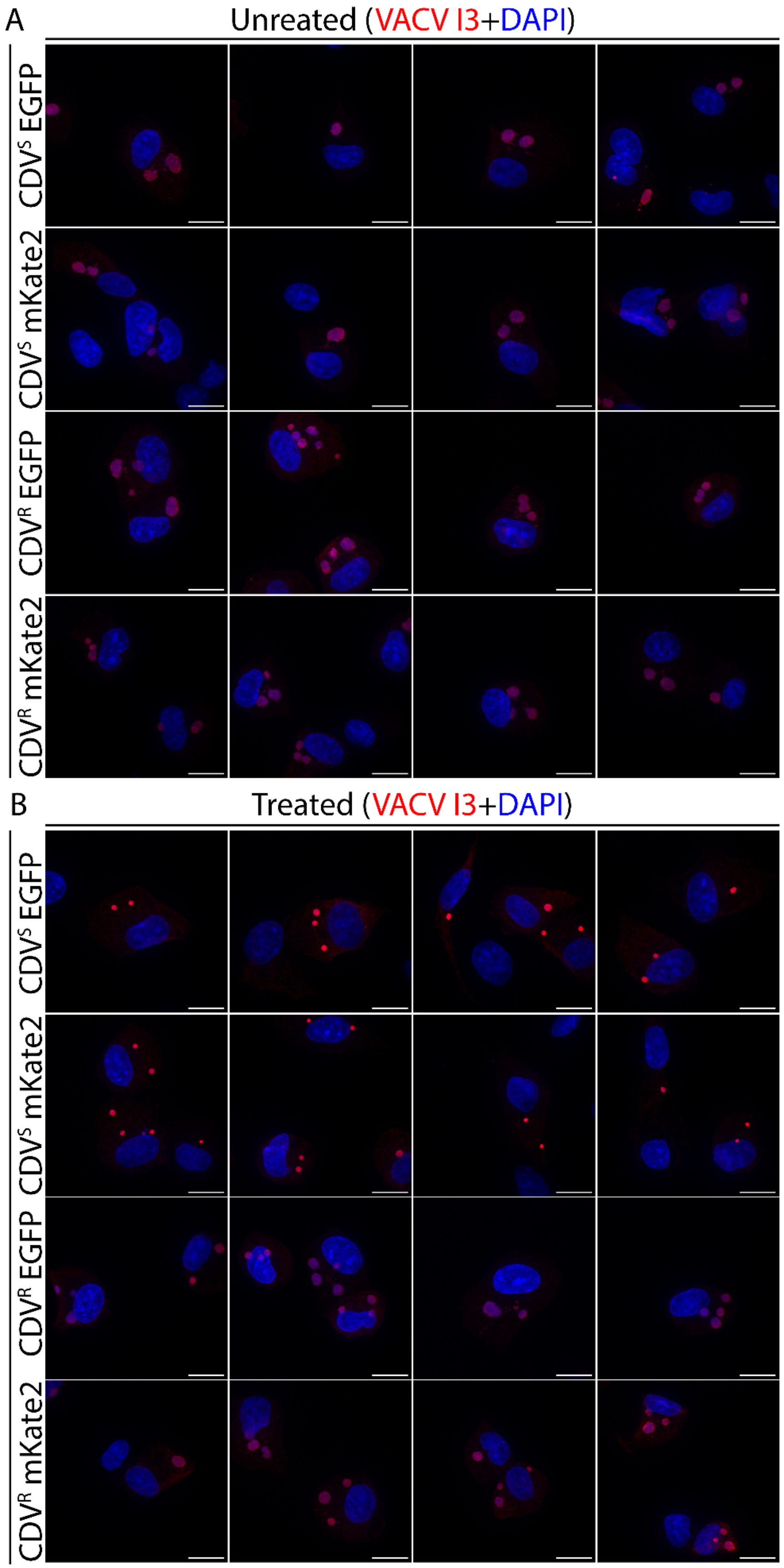
A
Unreated (VACV I3+DAPI)
CDV^S EGFP
CDV^S mKate2
CDV^R EGFP
CDV^R mKate2
B
Treated (VACV I3+DAPI)
CDV^S EGFP
CDV^S mKate2
CDV^R EGFP
CDV^R mKate2

**Supplemental Figure 3. Additional IF images of CDV$^S$ and CDV$^R$ factories.** The cells were infected and stained as in Figure 6A&B. Four fields of view are shown in parallel for each strain and treatment condition.

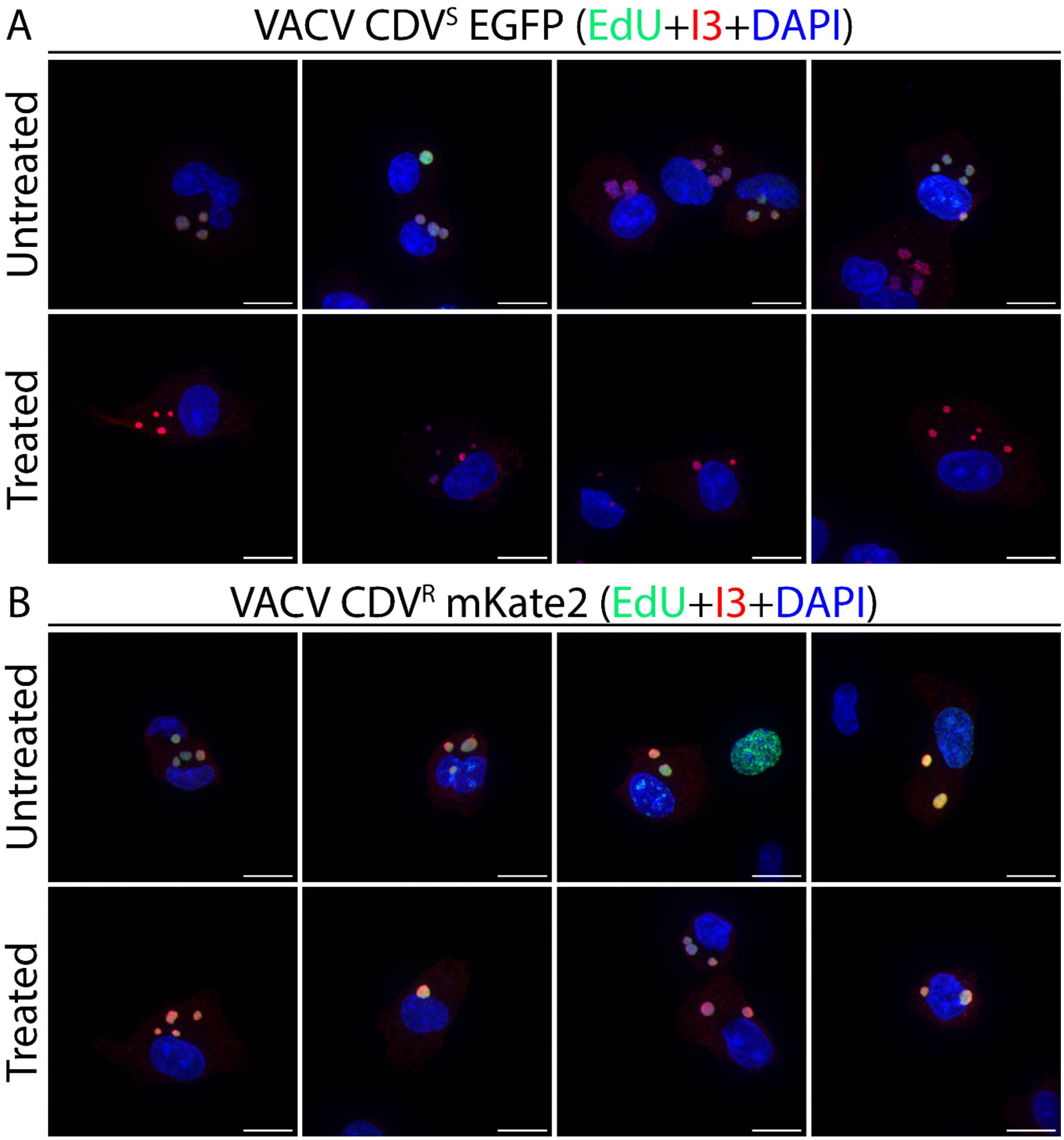


**Supplemental Figure 4. Additional IF images of EdU-labeled CDV$^S$ and CDV$^R$ virus factories.** The cells were infected and stained as in Figure 6D&E. Four fields of view are shown in parallel for each strain and treatment condition.

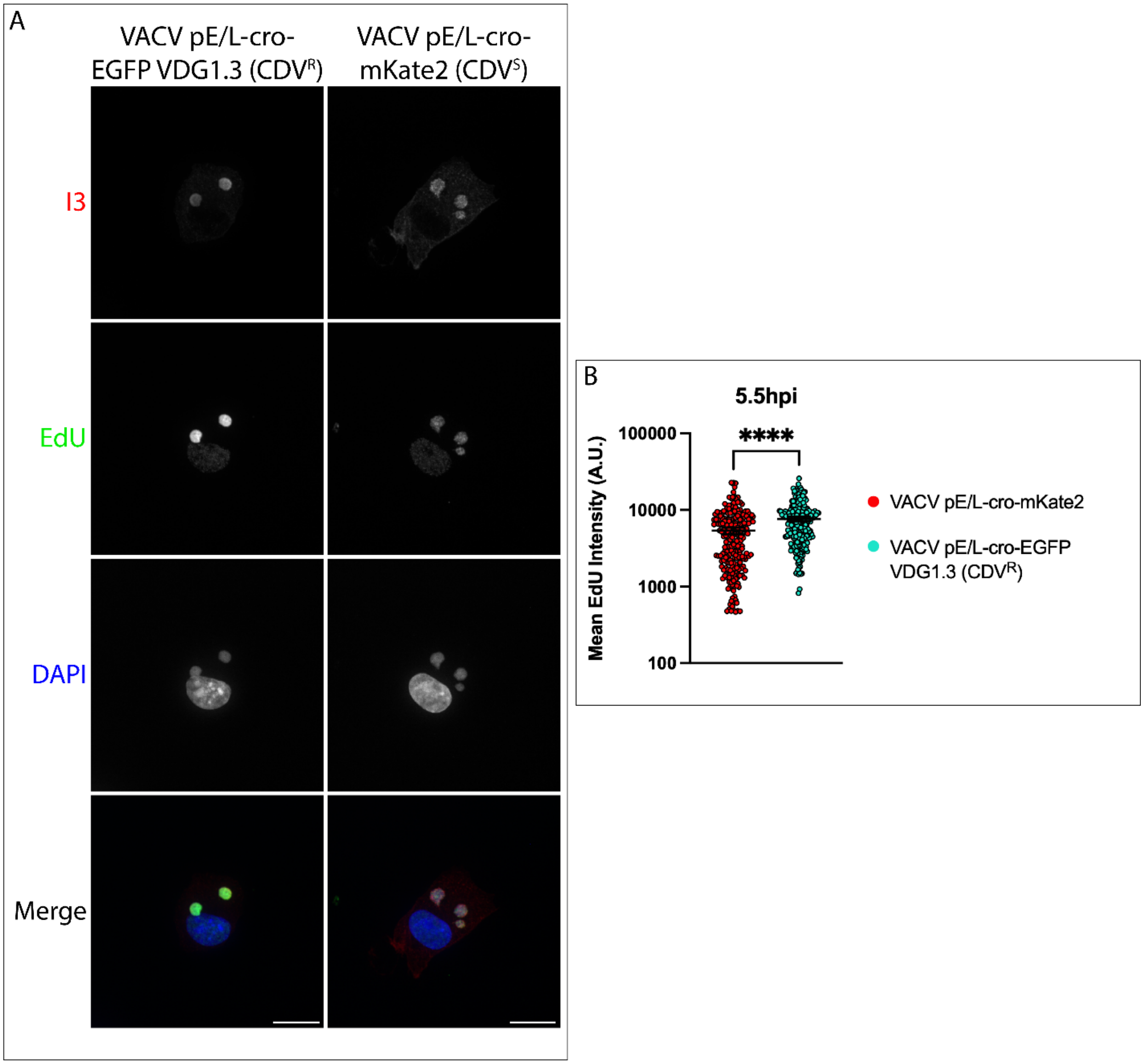


**Supplemental Figure 5. CDV$^R$ virus factories show higher levels of EdU incorporation than CDV$^S$ virus factories.** A) Representative images from cells infected with CDV$^R$ EGFP or CDV$^S$ mKate2 and pulsed with 10µM EdU at 5 hours post-infection. The cells were fixed at 5.5 hours post-infection and the EdU was labelled with Alexa Fluor 647 Azide by click chemistry. The I3 antibody and DAPI counterstain were used to visualize the virus factories. Scale bar = 15µm. B) Quantification of EdU-647 fluorescence intensity from 300 virus factories imaged across three independent experiments and analyzed using an unpaired, two-way t test. The data are plotted as the mean and 95% confidence interval.

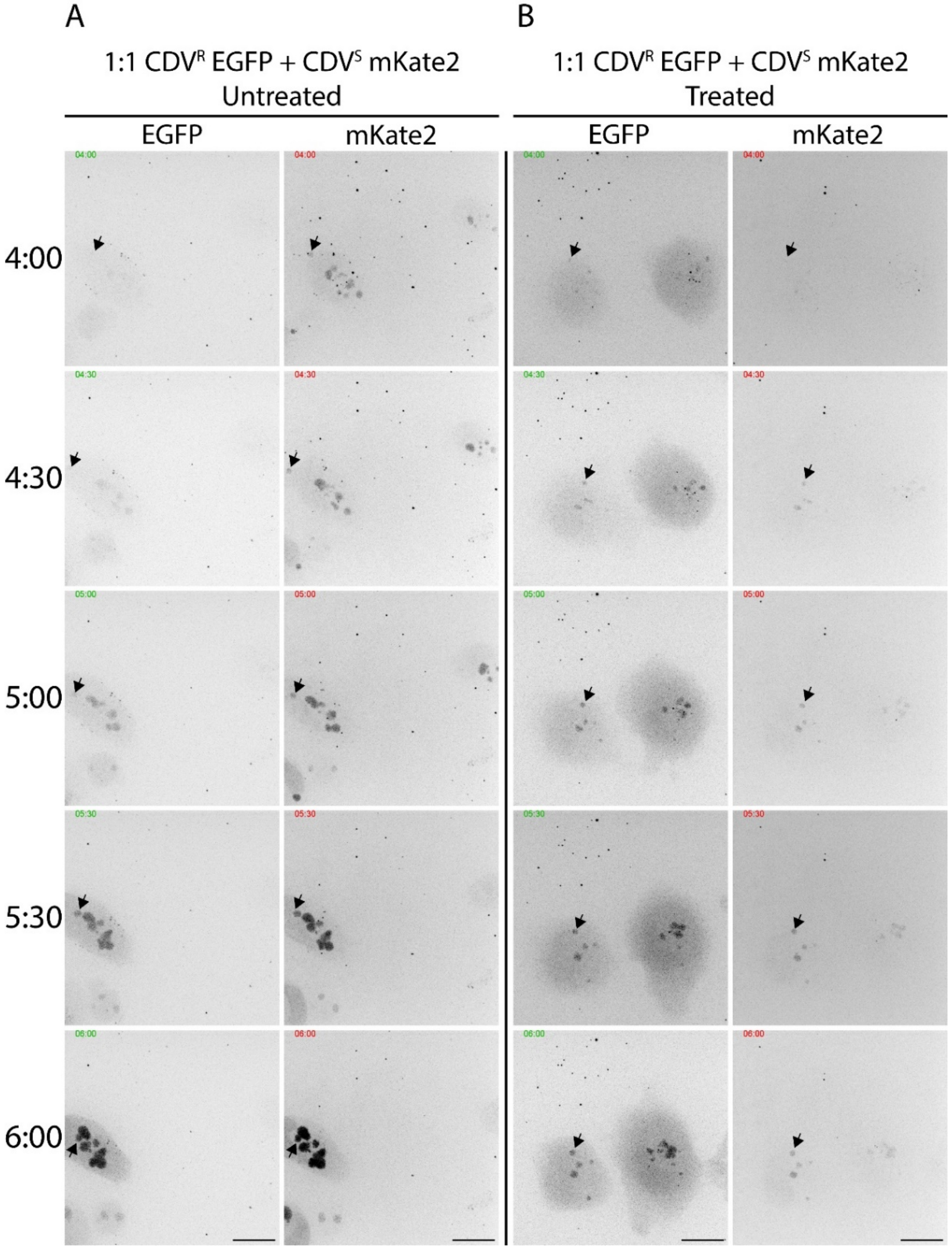


**Supplemental Figure 6. Live-cell imaging of cells coinfected with $CDV^R$ EGFP and $CDV^S$ mKate2.** BSC-40 cells were synchronously coinfected at a combined MOI of 5 with VACV $CDV^R$ EGFP and $CDV^S$ mKate2 in the absence (A) or presence (B) of CDV. The infections were imaged from 3-6 hours post-infection. Representative images from more than three independent biological experiments are shown, and the EGFP and mKate2 fluorescence for each field of view are shown side-by-side. Arrows indicate factories that did not fuse with adjacent factories throughout the experiment. Scale bar = 15μm.

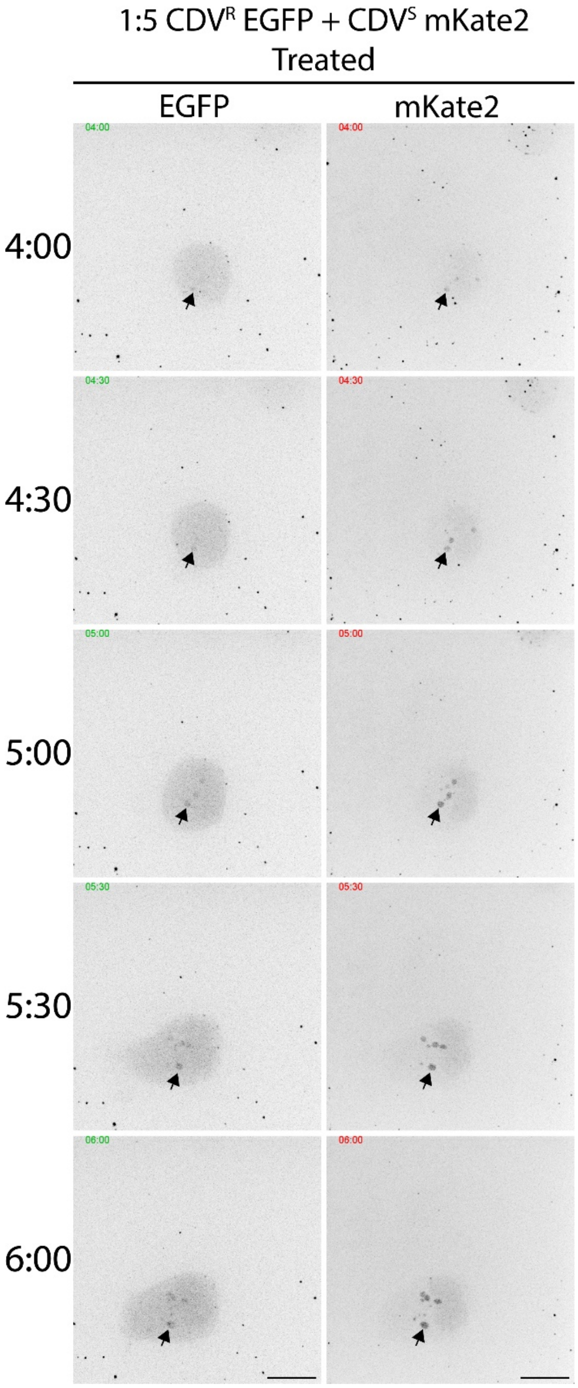


**Supplemental Figure 7. Live-cell imaging of coinfected cells.** BSC-40 cells were synchronously coinfected with VACV CDV$^{R}$ EGFP and CDV$^{S}$ mKate2. The combined MOI was 6, with a 1:5 ratio of CDV$^{R}$ and CDV$^{S}$ viruses. The infections were imaged from 3-6 hours post-infection. Representative images from more than three independent biological experiments are shown, and the EGFP and mKate2 fluorescence for each field of view are shown side-by-side.

Arrows indicate factories that did not fuse with adjacent factories throughout the experiment. Scale bar = 15µm.

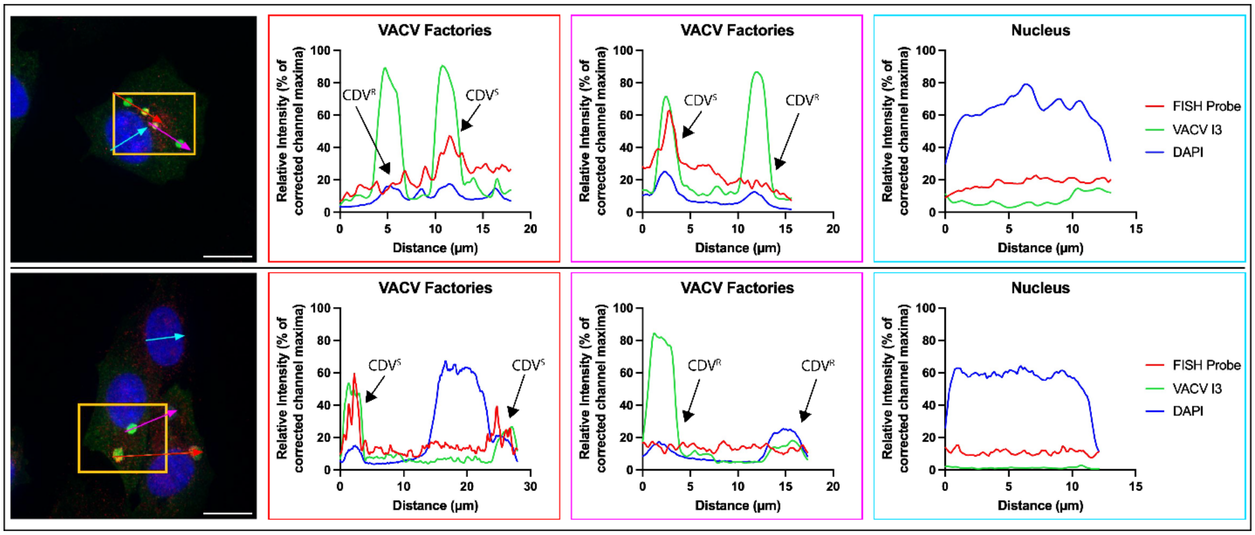


**Supplemental Figure 8. Line profiles of VACV infected cells used for FISH.** The images from Figure 7 are shown on the left. Scale bar = 15µm. The fluorescence intensities recorded in the individual channels were extracted from Volocity software and a background correction was applied based on the black limits for the entire image. Values were then normalized to the corrected channel maxima to create a relative fluorescence intensity for display purposes. The three graphs correspond to the different colored arrows and are coordinated by the colored boxes. Clear peaks in fluorescence intensity from the FISH probe and VACV I3 immunofluorescence were detected in some viral factories, which were designated as CDV$^S$ factories. The lack of strong peaks of fluorescence intensity in the nuclei indicate that the probe was not hybridizing to DNA lacking the target *gfp* gene.
